# Two determinants of dynamic adaptive learning for magnitudes and probabilities

**DOI:** 10.1101/2023.08.18.553813

**Authors:** Cedric Foucault, Florent Meyniel

## Abstract

Humans face a dynamic world that requires them to constantly update their knowledge. Each observation should influence their knowledge to a varying degree depending on whether it arises from a stochastic fluctuation or an environmental change. Thus, humans should dynamically adapt their learning rate based on each observation. Although crucial for characterizing the learning process, these dynamic adjustments have only been investigated empirically in magnitude learning. Another important type of learning is probability learning. The latter differs from the former in that individual observations are much less informative and a single one is insufficient to distinguish environmental changes from stochasticity. Do humans dynamically adapt their learning rate for probabilities? What determinants drive their dynamic adjustments in magnitude and probability learning? To answer these questions, we measured the subjects’ learning rate dynamics directly through real-time continuous reports during magnitude and probability learning. We found that subjects dynamically adapt their learning rate in both types of learning. After a change point, they increase their learning rate suddenly for magnitudes and prolongedly for probabilities. Their dynamics are driven differentially by two determinants: change-point probability, the main determinant for magnitudes, and prior uncertainty, the main determinant for probabilities. These results are fully in line with normative theory, both qualitatively and quantitatively. Overall, our findings demonstrate a remarkable human ability for dynamic adaptive learning under uncertainty, and guide studies of the neural mechanisms of learning, highlighting different determinants for magnitudes and probabilities.

**Significance statement:** In a dynamic world, we must constantly update our knowledge based on the observations we make. However, how much should we update our knowledge after each observation? Here, we have demonstrated two principles in humans that govern their updating and by which they are capable of dynamic adaptive learning. The first principle is that when they observe a highly surprising event indicating a likely change in the environment, humans reset their knowledge and perform one-shot learning. The second principle is that when their knowledge is more uncertain, humans update it more quickly. We further found that these two principles are differentially called upon in two key learning contexts that could be associated with different brain mechanisms: magnitude learning (which primarily requires adaptation to surprise, under the first principle) and probability learning (which primarily requires adaptation to uncertainty, under the second principle). Our findings advance understanding of the mechanisms of human learning, with implications for the brain and the development of adaptive machines.

## Introduction

Humans live in a dynamic world that requires them to constantly update their knowledge as events unfold. For example, new construction works that cause significant delays in transportation should prompt users to revise their estimate of their typical commute time. This learning process is a fundamental part of our adaptive behavior, and a field of study that spans the sciences of behavior, brain, and machines (Rescorla & Wagner, 1972; Rosenblatt, 1961; Schultz et al., 1997; Sutton & Barto, 2018).

Learning is challenging because the environment is often not only dynamic, but also stochastic. When we take public transportation, for example, our commute time varies from day to day due to stochastic fluctuations. However, a delay can also be caused by an abrupt change, such as construction work, that may persist for several weeks. In this case, we face an ambiguity when a delay occurs: is it due to stochastic fluctuations (in which case we should not significantly change our estimated average commute time), or a genuine change point (in which case we should revise our estimate more drastically)? The challenge lies in determining the appropriate weight to give to the current observation (the observed delay) in our learning process.

A descriptive tool to quantify the weight given by the learner to an observation is the *apparent learning rate*, which we will simply call *learning rate* hereafter (Heilbron & Meyniel, 2019; Nassar et al., 2010). The learning rate measures the amount of update of the learned value (from *v_t-1_* to *v_t_*) induced by the observation *x_t_* in proportion to its deviation from the previously learned value:

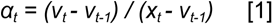

This tool is useful for characterizing the learning process and distinguishing several levels of adaptive learning (Foucault & Meyniel, 2021; Soltani & Izquierdo, 2019). Level zero (non-adaptive) corresponds to a constant learning rate, as in the delta rule, a widely used learning model in psychology and neuroscience. This model accounts for the basic fact that the amount of update increases with the discrepancy between the expectation and the observation (also known as prediction error). It has led to numerous successes across various forms of learning, including associative learning and reinforcement learning in humans, animals, and artificial agents (O’Doherty et al., 2003; Rescorla & Wagner, 1972; Rosenblatt, 1961; Schultz et al., 1997; Sutton & Barto, 2018). Beyond level zero, adaptive learning processes involve an adaptation of the learning rate at some level. We distinguish two levels of adaptive learning: one is at the average level over a block of trials, and the other is more fine-grained, at the level of individual trials.

A first level of adaptive learning corresponds to adjusting the average learning rate to the global statistics of the environment. Two relevant statistics for adjusting the average learning rate are the average frequency of change points (also known as volatility) and the level of stochasticity in the environment (Soltani & Izquierdo, 2019). To illustrate, change points are more frequent in transportation systems where construction campaigns are more frequent, and commute times are less stochastic in transportation systems that are better organized. When change points are more frequent, the average learning rate should be higher to update the estimated value more quickly. Accordingly, previous studies have shown that humans use a higher average learning rate in a block of trials containing many change points compared to a block with no change points (Behrens et al., 2007; Browning et al., 2015; Cook et al., 2019). When the environment is more stochastic, the average learning rate should be lower to stabilize the estimated value. Such an effect has also been observed in humans (Lee et al., 2020).

A second level of adaptive learning corresponds to dynamically adjusting the learning rate from one observation to the next depending on what is observed—we refer to it as *dynamic adaptive learning*. Such dynamic adjustments are particularly critical to learn effectively in a dynamic and stochastic environment, so as to increase the learning rate locally when a single change point is detected (Foucault & Meyniel, 2021; Nassar et al., 2010). Compared to the first level (different average learning rates between blocks of trials), less is known in humans about this second level (dynamic adjustments of the learning rate at the trial level).

Dynamic adaptive learning has been demonstrated in humans in the case of magnitude learning. Empirical evidence is available for several kinds of magnitudes including symbolic numbers (Nassar et al., 2012), positions (McGuire et al., 2014), and angular directions (Vaghi et al., 2017). However, these studies leave unexplored another common type of learning: probability learning. In the transportation example, this could be the probability that a traffic jam (or an incident in transit) occurs and causes us to be late.

The literature on probability learning is vast, from the learning of reward probabilities to stimulus occurrence to the validity of attentional cues, but it remains unknown whether humans dynamically adapt their learning of probabilities. Previous studies are not suitable to examine trial-level adjustments of the learning rate. Many of them are based on choices, from which only an average learning rate can be estimated, as a model parameter that is fitted across many choices and compared between blocks of trials (Behrens et al., 2007; Browning et al., 2015; Cook et al., 2019; Cools et al., 2002).

From a learning perspective, magnitudes and probabilities present a fundamental difference: the amount of information provided by a single observation about the quantity to be learned is typically much lower for probabilities than for magnitudes (see Fig. S1 for common examples). This amount of information should in principle regulate the learning rate: the more information the observation provides about the quantity to be learned, the higher the learning rate should be. In particular, a single observation may be informative enough to detect a change point and thus immediately increase the learning rate. Conversely, when observations are less informative, change points are more ambiguous (difficult to distinguish from stochastic fluctuations), making dynamic adaptive learning all the more challenging.

In light of these differences, what has been demonstrated about human adaptive learning in the context of magnitudes may not generalize to probabilities. A comprehensive understanding of dynamic adaptive learning requires to study probability learning and to compare it directly to magnitude learning, in order to uncover the extent and determinants of dynamic adjustments in each case. Here, we address two main questions:

1. Do humans dynamically adapt their learning rate in probability learning (level 2, introduced above) or use a fixed learning rate (level 0)? What are their adjustment dynamics, and how do they compare to those of magnitude learning?
2. What are the computational determinants of the dynamic adjustments of learning rates?

To answer these questions, we conducted a study in which human subjects performed two learning tasks, a magnitude learning task and a probability learning task. We developed a new experimental paradigm to measure the subject’s learning rate directly from their report at each observation. This measure of the learning rate serves to characterize the human learning process.

To identify the determinants of adjustments in learning rates, we adopted a normative approach (also known as rational analysis) (Lieder & Griffiths, 2020; Oaksford & Chater, 2007). We studied the optimal Bayesian solution to the learning problem posed and derived normative theoretical properties, which we then tested in subjects. We want to stress that we make no claims about the algorithm used by subjects to achieve these properties (we use the Bayesian solution as a means to identify these properties, not as a model of the algorithm).

We identified two computational determinants and we show that they play a quantitatively different role in the two types of learning. Overall, our results show a remarkable alignment of human behavior with normative behavior in dynamic adaptive learning, even in the difficult case of probability learning.

## Results

### Measuring the learning rate at each observation during magnitude or probability learning

To measure subjects’ learning rate at each observation as they are learning, we developed a new experimental paradigm where subjects update their estimate in real time as they observe a sequence of stimuli occurring one at a time at regular intervals of 1.5s (Fig. 1). We obtained real time reports by continuous motion tracking: the estimation is done with a slider that follows the subject’s motion, captured on a touchpad or a mouse (Fig. 1 A and B). The subject’s estimate (*v_t_*, obtained from the slider position) is thus collected at each observation (*x_t_*, *t* indexes the observation). This allows us to calculate the learning rate for each observation using equation [1].

**Fig. 1.**
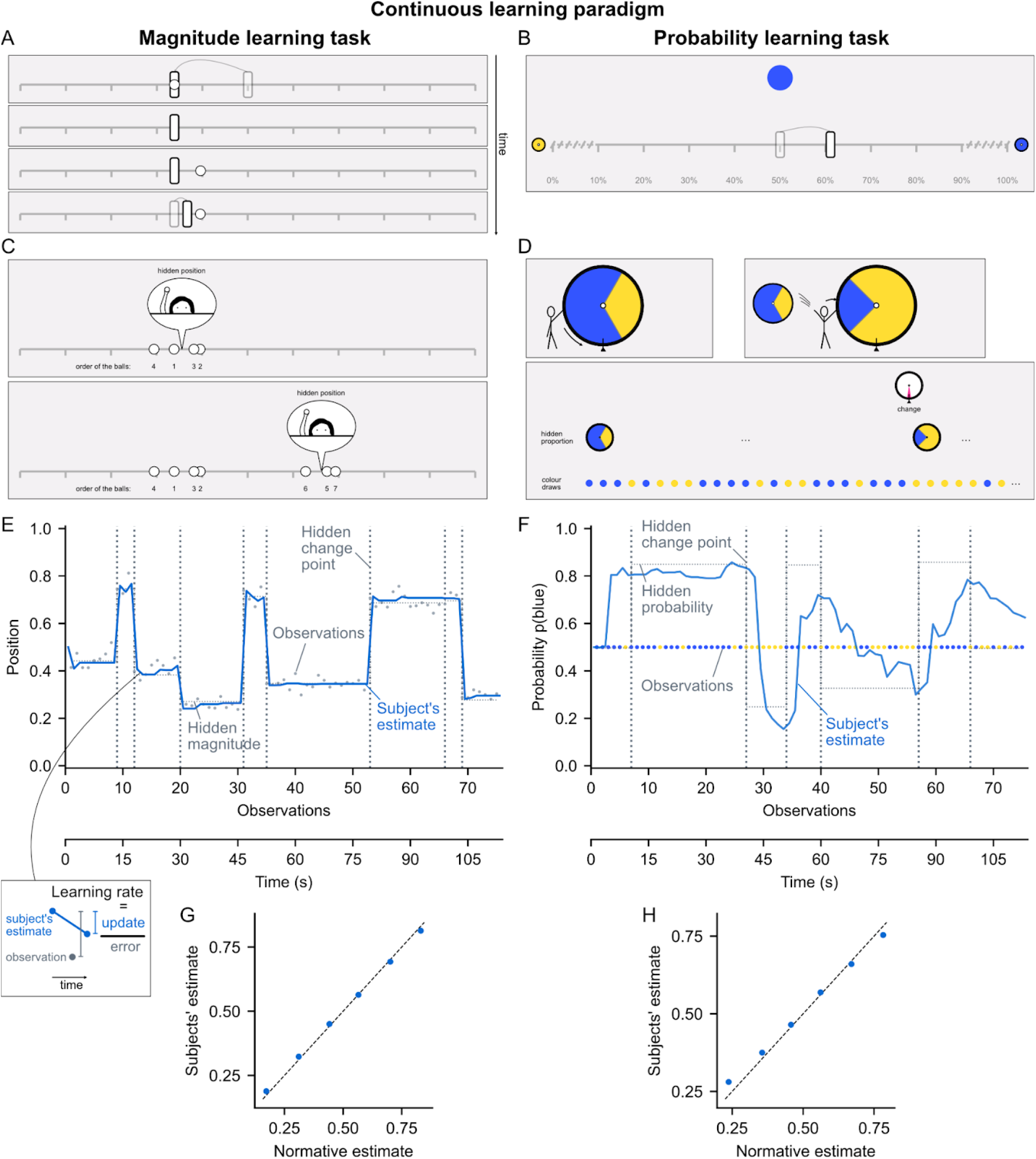
Magnitude and probability learning tasks with estimates reported in real-time. (A and B) Screenshots of the two tasks. The subject must estimate a hidden quantity (magnitude or probability) based on the stimuli which appear one by one at regular intervals (1.5s). This quantity changes at hidden, unpredictable times called change points. The subject reports their current estimate by moving a slider in real time via motion tracking. For the magnitude task (A, frames show successive times), the hidden quantity is the mean horizontal position generating the stimuli (white circles). For the probability task (B), it is the probability that the circle appearing in the center is blue vs. yellow. Semi-transparent elements indicating slider movements were added here for explanation. (C and D) Screenshots of the task instructions. For the magnitude task (C), subjects were told that they should estimate the position of a hidden person throwing snowballs based on where the balls are landing (the white circles), and that the person could change position at random times without their knowledge. For the probability task (D), subjects were told they should estimate the proportion of blue and yellow on a hidden wheel used to draw the observed colors, and that the wheel could be changed at random times without their knowledge. (E and F) Example of one session of the magnitude (E) and probability (F) task. Vertical dotted lines indicate the first observation after a change point. Thin dotted lines show the hidden quantity values. Dots represent observations (positions in (E), colors in (F)). The blue line shows the subject’s estimate after each observation (0: left edge of the slider, 1: right edge). (G and H) Accuracy of subjects’ estimates for each observation compared to normative (i.e. optimal) estimates in the magnitude (G) and probability (H) task. The data were binned in 6 equal quantiles of normative estimate and averaged within-subjects. Points and error bars (too small to be seen) show mean ± s.e.m. across subjects, dashed line is the identity line.

Note that the learning rate calculated in this way is directly obtained from the subject’s estimates (Nassar et al., 2010), rather than derived from model fitting, as is often done (Behrens et al., 2007; Browning et al., 2015; Cook et al., 2019). This learning rate is a behavioral measure that makes no assumptions about the computational process used by subjects for learning. The learning rate can thus be measured for all kinds of learning models and compared to subjects.

We developed two learning tasks using this paradigm: a magnitude learning task and a probability learning task. In order to compare their results, we adopted a similar design and task structure. In both tasks, the subject’s goal is to estimate a hidden quantity (magnitude or probability) from the observations they receive, and to produce estimates as accurate as possible at each observation. During the observation sequence, the hidden quantity undergoes discrete changes at unpredictable random times called change points (not limited to reversals), which are also hidden (subjects are not told when a change point occurs). In the magnitude learning task, the hidden magnitude is the mean horizontal position generating the observed stimulus positions. Subjects were instructed that a hidden person was throwing snowballs and that their goal was to estimate the position of this hidden person based on where the snowballs are landing, knowing that the hidden person could change position from time to time without their knowledge (Fig. 1C). In the probability learning task, the hidden probability is the probability of the centrally-presented stimulus being blue (vs. yellow). Subjects were instructed that the colors were drawn by spinning a wheel filled with a certain proportion of blue and yellow and that their goal was to estimate this proportion, knowing that the wheel could be changed from time to time without being told when it changes (Fig. 1D). The observation sequences were generated, in the magnitude task, using the same parameters as (Nassar et al., 2012) for comparison (same or very similar parameters were also used in other magnitude learning studies (McGuire et al., 2014; Vaghi et al., 2017)). In the probability task, we used a probability of change point occurrence that made the task engaging and difficult but still tractable for subjects. (See Methods for more details.)

The human subjects (n=96) performed the two task one after the other, in a counterbalanced order across subjects (see an example session for each task Fig. 1 E and F). We evaluated subjects’ estimate accuracy at the level of each observation by comparing their estimates with the normative estimates, i.e. the optimal estimates given the observations received. Although change points were frequent, subjects were able to produce accurate estimates (close to normative) of magnitudes and probabilities (Pearson *r*=0.96±0.01 and 0.80±0.01 mean±s.e.m., t_95_>55.7, p<10^-73^ in both cases, see Fig 1 G and H). (See Supplementary Text 1 for further details on the comparison between the normative estimates and the subjects’ estimates.)

### Subjects dynamically adapt their learning rate after a change point for probabilities, and differently than for magnitudes

As per question 1), we investigated subjects’ ability to dynamically adapt their learning rate in response to change points, by calculating their learning rate with equation [1] and analyzing its dynamics around the change points. An optimal learner should increase their learning rate when they believe that a change point has occurred recently, to quickly update their estimate, and then decrease their learning rate to stabilize their estimate (see illustration of this behavior and how it differs from that of a learner using a fixed learning rate in Fig. 2C). Note that change points are more difficult to detect in the probability learning task because the amount of information provided by a single observation is much lower than for magnitude learning (Fig. S1). Despite this difficulty, subjects dynamically adapted their learning rate remarkably well, not only in the magnitude learning task, but crucially also in the probability learning task (Fig. 2 A and B).

**Fig. 2.**
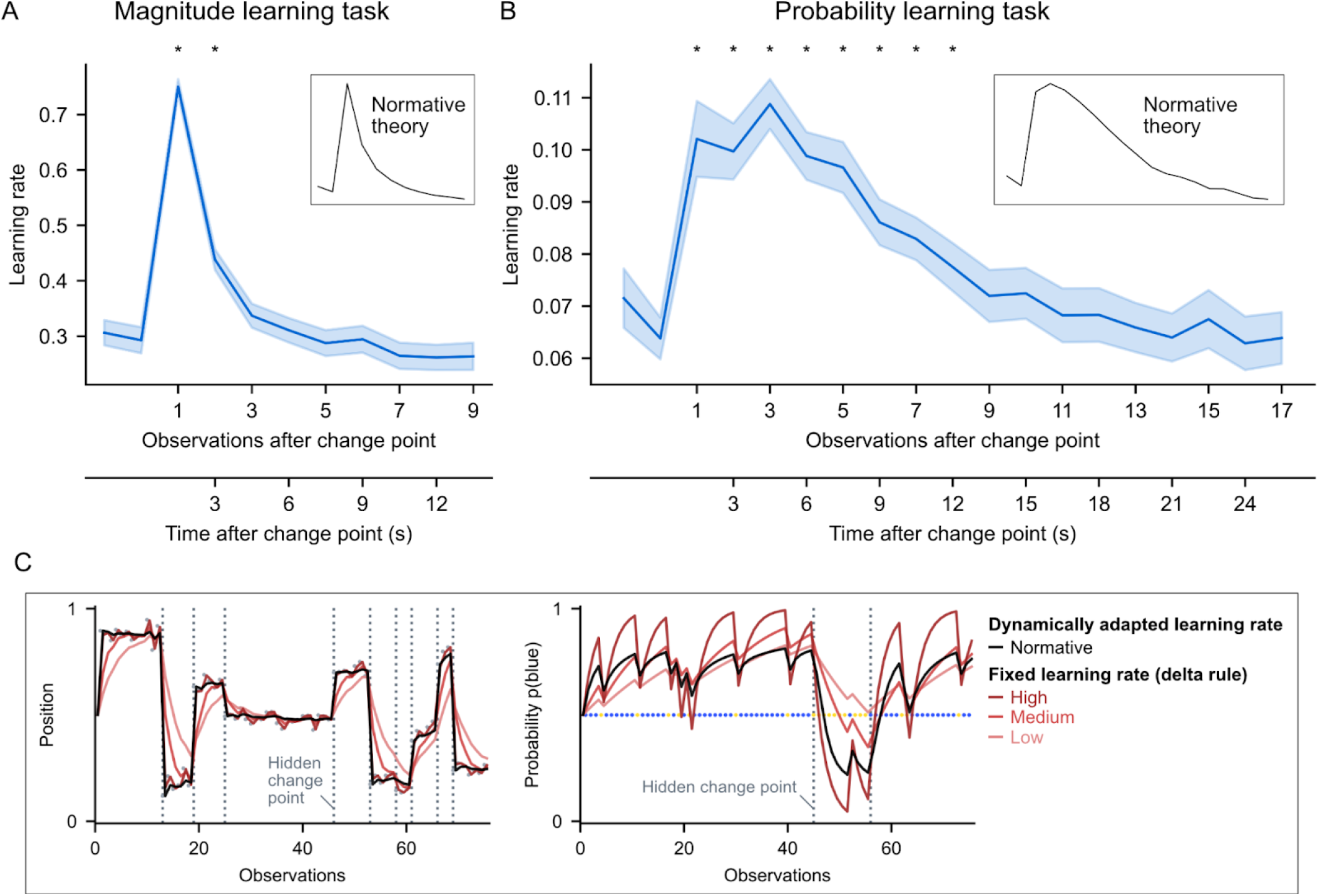
Dynamics of subjects’ learning rate in response to a change point in the two tasks. The subjects’ learning rates were measured at each observation (equation [1]), then aligned to change points and averaged within-subject. Lines and error bands show mean ± s.e.m. across subjects. Subjects dynamically adapt their learning rate after a change point, increasing it transiently and mainly at the first observation in the magnitude learning task (A), and more prolongedly in the probability learning task (B). Stars show statistically significant differences compared to the baseline measured at the two observations before the change point (p<0.05, two-tailed, FWE cluster-corrected for multiple comparisons across time). The subjects’ learning rate dynamics are very similar to those prescribed by normative theory, shown in the insets. (C) Examples illustrating the difference in behavior between a fixed learning rate (delta-rule, red curves, for different values of the learning rate) and a dynamic one (normative learner, black curve). When the learning rate is fixed, regardless of its value, the behavior shows systematic deviations making learning less efficient: after a change point, the estimates are too slowly updated, and in the absence of change points, they fluctuate too much following the random variability in observations. By dynamically increasing and then decreasing the learning rate after a change point, the normative learner quickly updates and then stabilizes its estimate.

In the magnitude learning task, the learning rate dynamics we found (Fig. 2A) replicate those found in previous studies of magnitude learning (McGuire et al., 2014; Nassar et al., 2012; Vaghi et al., 2017). The increase in learning rate occurs immediately and mainly at the first observation after a change point, with a learning rate suddenly close to 1. In the probability learning task, the learning rate dynamics show an increase that is smoother and more prolonged, remaining significantly higher at the eighth observation after a change point compared to before the change point (Fig. 2B). Importantly, these different dynamics are very similar to those prescribed by normative theory (see insets in Fig. 2; the Pearson correlation between subject’s mean dynamics and the normative one is 0.99 in both tasks; the amplitudes of their dynamics are also similar to the normative ones, shown in further detail in Fig. S2). We also controlled for the frequency at which subjects updated: we observed similar, significant dynamic adaptations of the learning rate even after excluding from analysis all observations where subjects did not report an update (Fig. S4). (Note that subjects frequently updated their estimate in our study, see Fig. S3).

### Two normative determinants of the learning rate

Having demonstrated the dynamic nature of subjects’ learning rate, we sought to identify the determinants that drive these dynamics (question 2). To this end, we formalized the two tasks within the same probabilistic framework and resorted to the general principles that govern learning according to normative theory. Below, we summarize the essential elements for understanding the normative determinants identified.

According to normative theory, optimal estimates are obtained by calculating the posterior distribution *p(h_t_ | x_1:t_)*, that is, the posterior distribution for the value of the hidden quantity (*h_t_*), given the observations received so far in the sequence (*x_1:t_*). The posterior must be updated each time a new observation is received. The normative updating process, whereby the new posterior is calculated from the previous one and the last observation, can be broken down into two steps (see Fig. 3A and equation [4] in Methods). In the first step, the previous posterior (that is, the posterior on *h_t-1_*) is transformed into a prior on *h_t_* by incorporating the generative probability of a change point occurring between the two. In the second step, this prior is multiplied with the likelihood function of the last observation to give the new posterior (that is, the posterior on *h_t_*).

**Fig. 3.**
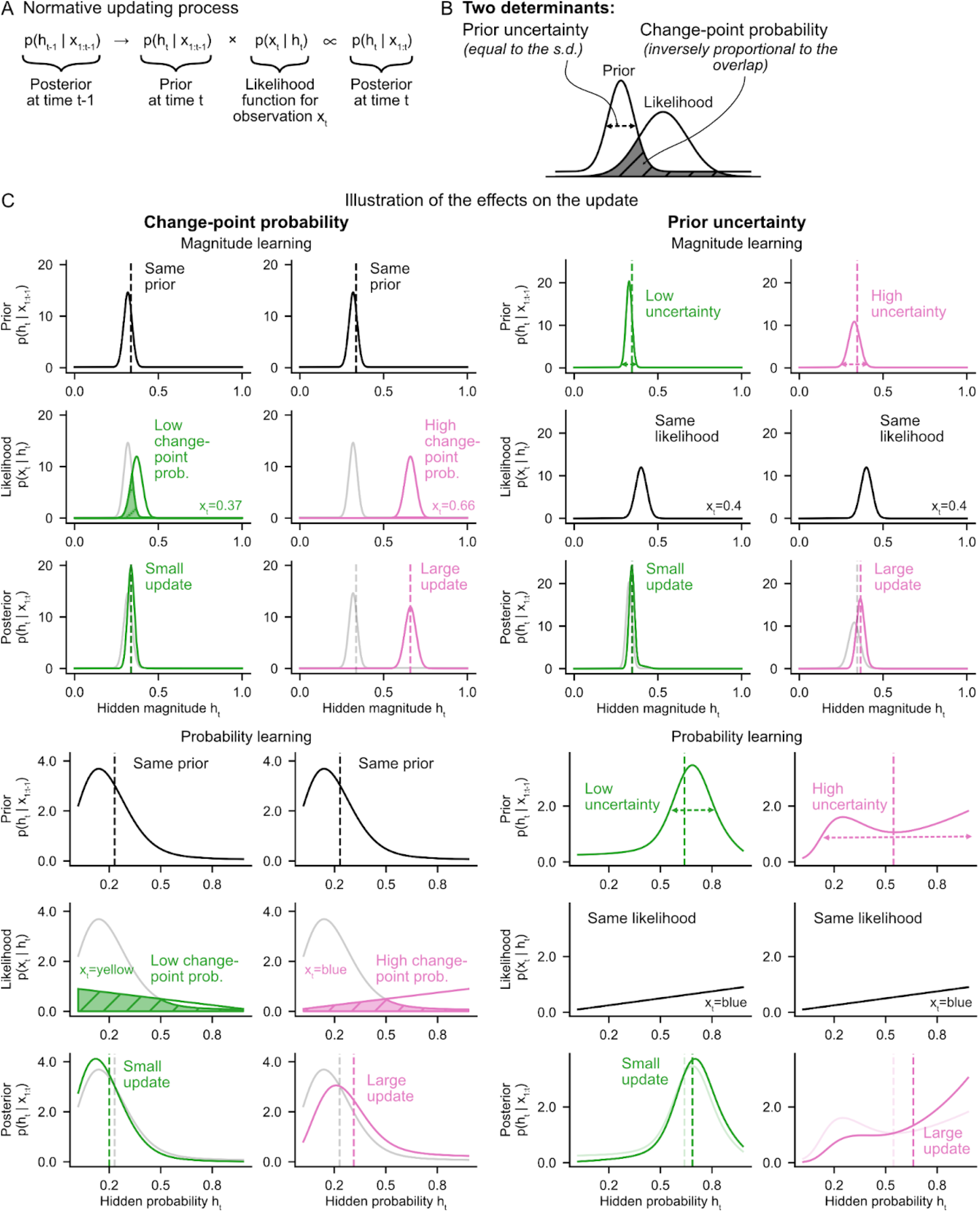
Two determinants regulate updating in normative theory. (A) The normative updating process involves the product of the prior, *p(h_t_|x_1:t-1_)*, and the likelihood of the last observation, *p(x_t_|h_t_)*. (B) The amount of update resulting from the product is determined by two factors: the uncertainty of the prior (dispersion), and the change-point probability indicated by the last observation (visualized by the degree to which the likelihood overlaps with the prior: the less it overlaps, the greater the change-point probability). (C) Illustration of the effect on the update of each determinant in each task. The specific effect of each determinant is shown by manipulating either the likelihood (through the last observation) or the prior (through the observations prior to last). A higher change-point probability or prior uncertainty each produce a greater update (visible by the shift of the distribution and its mean between the prior and posterior). Vertical dashed lines on the plots of the distributions indicate their means. On the likelihood and posterior plots, the prior is shown in semi-transparency to visualize the overlap and the update, respectively.

Two factors determine the amount of update (and therefore, the learning rate, which is proportional to the amount of update, equation [1]) that results from the product of the prior and the likelihood function (Fig. 3B).

i. The *prior uncertainty*, *u_t_*, quantified by the standard deviation of the previous posterior:

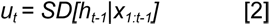

This is the uncertainty about the value of the hidden quantity before receiving the last observation. The greater the uncertainty, the larger the resulting update for the same observation, as the wider spread of the distribution leads to a larger shift (towards values where the likelihood is greater) when it is multiplied with the likelihood function. See Fig. 3C, right.

(ii) The *change-point probability*, *Ω_t_*, which is the term used in previous literature for the estimated probability that a change point occurred at the last observation (McGuire et al., 2014; Nassar et al., 2010, 2012; Vaghi et al., 2017). It is expressed as:

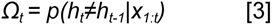

The change-point probability can be understood as the degree to which the likelihood function deviates from the prior. Visually, it corresponds to the degree to which the likelihood function and the prior overlap (Fig. 3B). The change-point probability is inversely proportional to this overlap (see denominator of equation [5] in Methods: the integral is equivalent to the area under the curve of the product of the two curves, which is strongly related to their area of overlap). It is also strongly related to the magnitude of the error elicited by the last observation, *x_t_ -v_t-1_*, also known as prediction error (Spearman correlation ρ>0.97 in each of the two tasks). A higher change-point probability leads to a larger update because, as the change-point probability increases, the previous posterior is increasingly discarded (by multiplication with the likelihood function). See Fig. 3C, left.

### Differential effect of the two determinants in magnitude and probability learning

Normative theory predicts a differential effect of change-point probability and prior uncertainty on the learning rate in the two tasks: the effect of change-point probability should dominate over that of prior uncertainty in the magnitude learning task, and the effect of prior uncertainty should dominate over that of change-point probability in the probability learning task (Fig. 4A). The effect sizes are measured by the weights of a multiple linear regression on the learning rate, and made comparable through z-scoring prior to regression. This differential effect is due to the fact that the variations of the two factors are not equal in the two tasks: change-point probability has much larger variations in magnitude learning compared to the probability learning, and conversely, prior uncertainty has much larger variations in probability learning compared to the magnitude learning (Fig. 4B). Thus, while both factors modulate the learning rate in both tasks, the factor that varies the most in the task is most responsible for the variations in the learning rate. The differences in variations of the two factors are related to differences in the likelihood function involved in magnitude vs. probability learning, which, in the case of magnitude, is more precise (see Fig. 3C, top vs. bottom), provides more information (Fig. S1), and thus significantly reduces uncertainty and can produce extreme values of change-point probability.

**Fig. 4.**
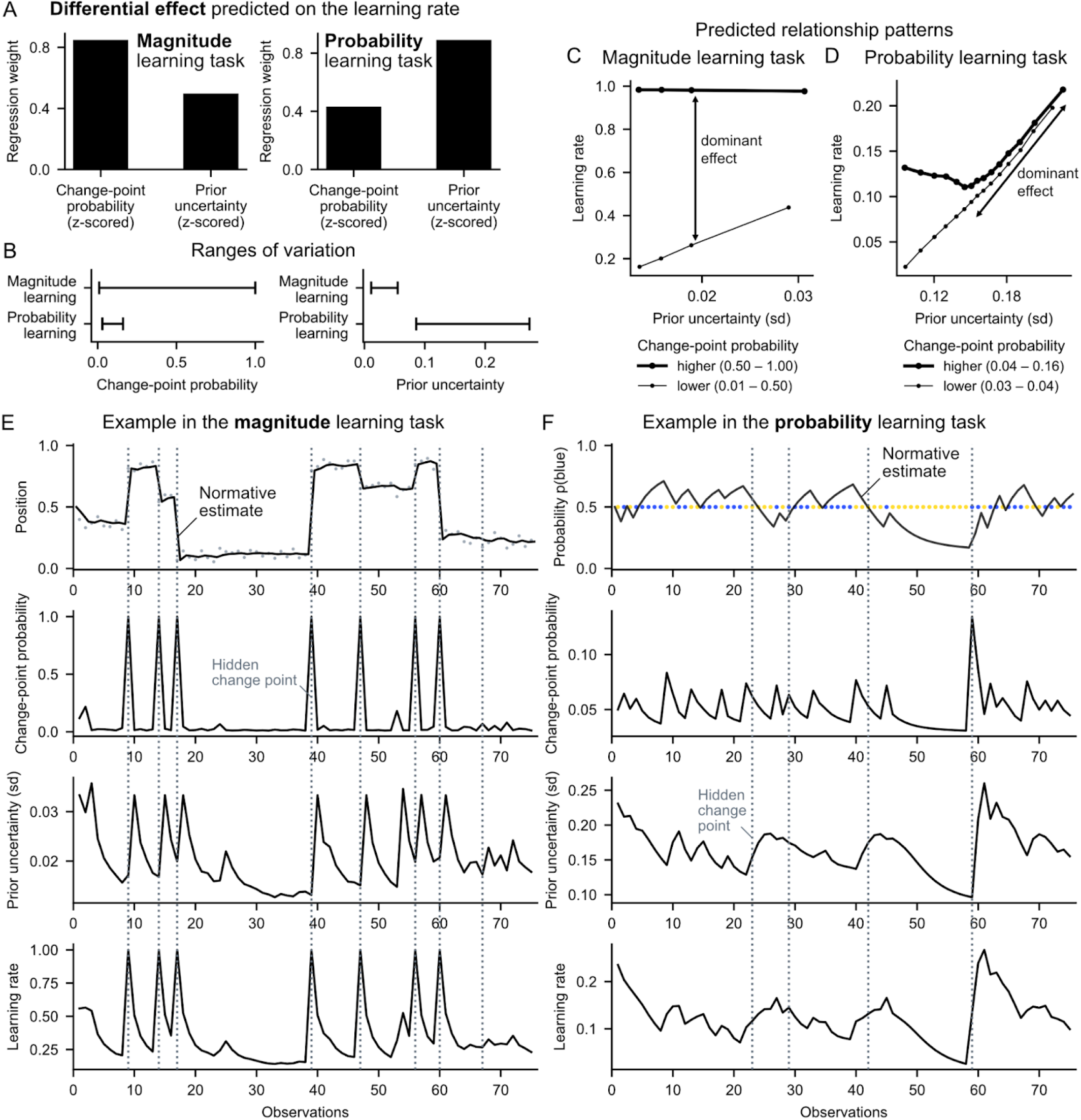
Differential effect predicted by normative theory in the two tasks. (A) In the magnitude task, the effect of change-point probability on the learning rate should dominate over that of prior uncertainty, whereas in the probability task, the effect of prior uncertainty should dominate. Effects are quantified by the weights of a multiple regression of the two factors on the normative learning rate (to make the weights commensurable, all variables were z-scored). (B) Ranges of variation (min, max) of the two determinants in the two tasks. (C and D) Relationship patterns predicted between the learning rate, prior uncertainty and change-point probability in the magnitude (C) and probability (D) task. In A, C and D, we carried out the analyses on the model in the same way as for subjects in Fig. 5. (E and F) Examples illustrating the dynamics of the different normative variables as a function of the sequence in the magnitude (E) and probability (F) task; note that the y-axis greatly differs between tasks for change-point probability, prior uncertainty and learning rate.

We further characterized the effects of change-point probability and prior uncertainty by relating them jointly to the learning rate and plotting the qualitative patterns that should be found in the two tasks (Fig. 4 C and D). These effects are best understood when viewed over time, by relating the variables to each other and to the observations, see Fig. 4 E and F.

Do human subjects regulate their learning rate according to the two determinant factors as predicted by normative theory? To find out, we performed the same analyses as we had done with the normative model but using the subjects’ learning rates instead. All the normative signatures we had identified were found in the subjects. First, the weights of change-probability and prior uncertainty on subjects’ learning rates are significant in both tasks (Fig. 5 A and B) (in the magnitude task, mean±s.e.m weight and two-tailed t-test against zero for change-point probability: 0.43±0.02, t_95_=23.4, p=10^-41^, and for prior uncertainty: 0.14±0.01, t_95_=13.8, p=10^-24^; in the probability task, for change-point-probability: 0.07±0.02, t_95_=4.3, p=10^-5^, and for prior uncertainty: 0.15±0.01, t_95_=17.0, p=10^-30^). Second, in the magnitude task, the weight of change-point probability is significantly higher than that of prior uncertainty (Fig. 5A; two-tailed paired samples t-test: t_95_=17.1, p=10^-30^; see also results in Supplementary Text 2 for comparison with previous studies). Conversely, in the probability task, the weight of prior uncertainty is significantly higher than that of change-point probability (Fig. 5B; t_95_=-4.4, p=10^-5^). Third, a graded co-variation of the learning rate with prior uncertainty can be observed in the magnitude task (Fig. 5C) and even more so in the probability task (Fig. 5D). Fourth, the interaction effects between prior uncertainty and change-point probability on the learning rate in subjects are similar to the normative ones (Fig. 5 E and F).

**Fig. 5.**
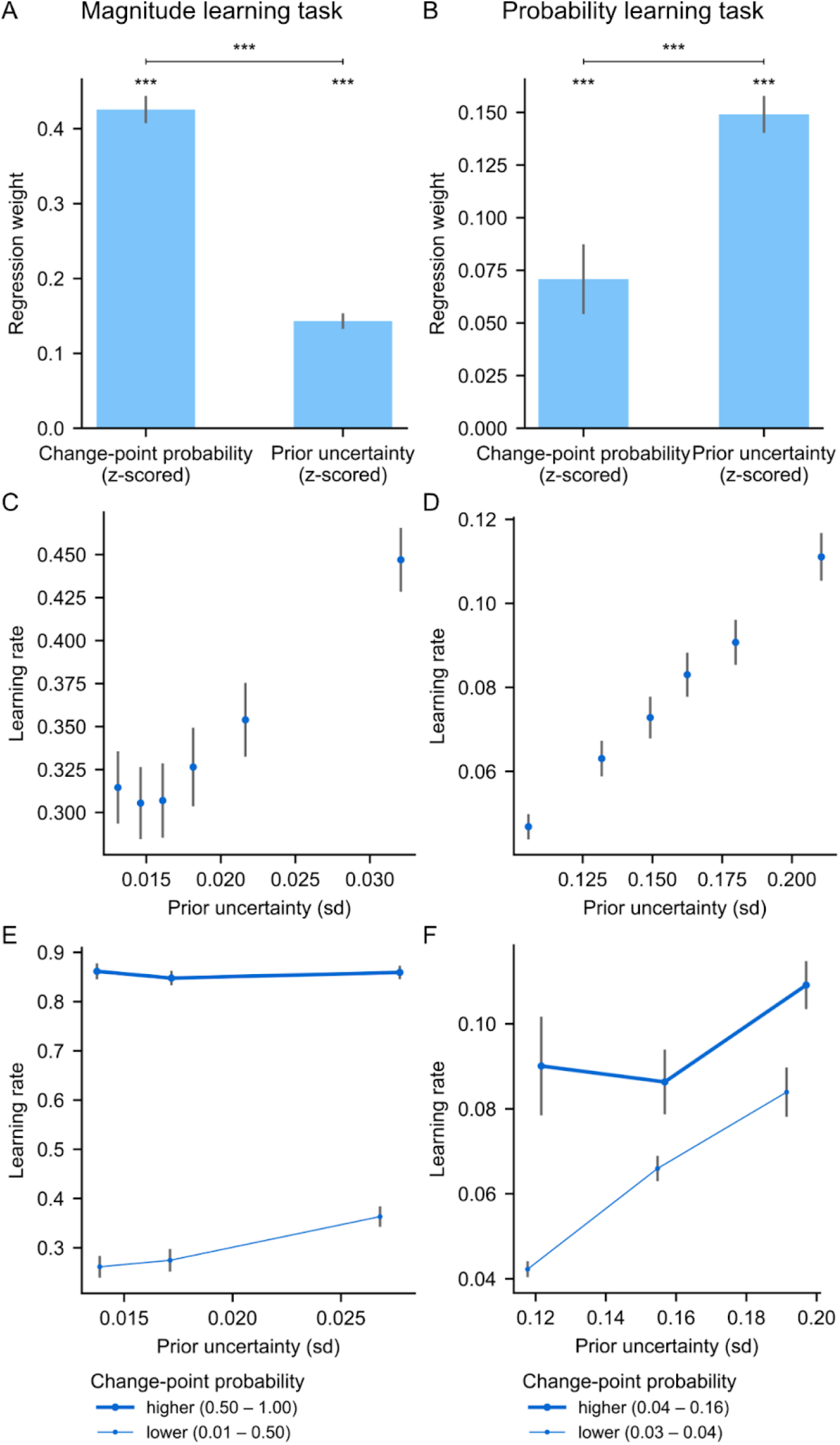
Subjects’ learning rates are differentially affected by the two determinants, as predicted by normative theory. (A and B) Weights of the change-point probability and prior uncertainty on subjects’ learning rates in the magnitude (A) and probability (B) task, calculated per subject by multiple regression on the subject’s learning rate. As predicted in Fig. 4A, the effect of change-point probability dominates in the magnitude task; that of uncertainty dominates in the probability task. ***: p<0.001, two-tailed t-tests. (C and D) Subjects’ learning rate increases with prior uncertainty, especially in the probability task. (E and F) The interaction effects between prior uncertainty and change-point probability on subjects’ learning rate are similar to the normative model (Fig. 4 C and D). In all plots, subjects’ learning rates were measured at each observation (equation [1]), and prior uncertainty and change-point probability were calculated using the normative model for the same observations. Bars/dots and error bars show mean ± s.e.m. across subjects.

### Average human learning is close to normative

So far, we have investigated the many ways in which human learning behaves normatively, but it might also have systematic deviations. Here, we studied systematicity at the group level by quantifying a systematic deviation as the degree to which the average estimate of human participants for a given observation deviates from normative estimate. Systematic deviations should be distinguished from non-systematic deviations, arising from mere variability. Both contribute to the deviations observed in subjects, but what proportion of those deviations can be attributed to the systematic vs. non-systematic component at the group level?

We disentangled the two components by performing a decomposition of the mean squared error. According to statistical theory, the mean squared error (*mse*) is the sum of two sources of error: the (squared) *bias error* (*sbe*), which is the systematic component, and the *variance error* (var), which is the non-systematic component (Hastie et al., 2009).

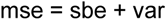

Here, we are interested in the error between the normative estimate and the subjects’ estimate for a given observation of the same sequence. To measure the bias and variance errors empirically in our study, we presented identical observation sequences across subjects (see Fig. 6, examples at the bottom). This allowed us to empirically calculate the mean and variance of the estimate across subjects, and the resulting bias and variance errors, for each observation of each sequence (see Methods for the mathematical expressions of this calculation). We then summed the errors across observations and sequences to obtain the proportion of the total mean squared error due to bias vs. variance. (Note that the variance measured in this way at the group level encompasses both inter-subject variability, which accounts for individual biases non-reproducible across individuals, and intra-subject variability as in (Drugowitsch et al., 2016), which is reflected across subjects in our case as there is at most one presentation per subject.)

**Fig. 6.**
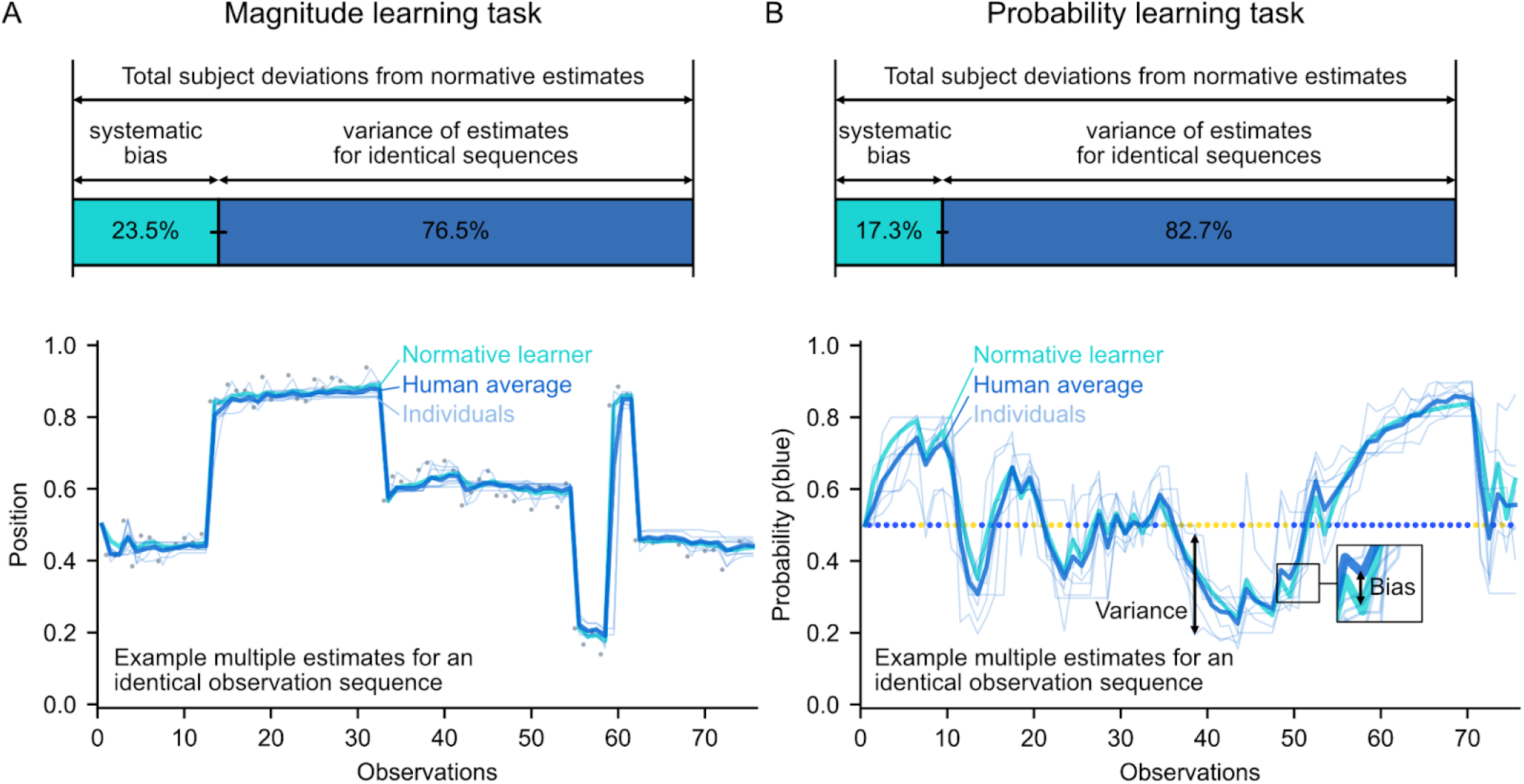
Group-level decomposition of deviations from normative learning reveals minor systematic deviations compared with the overall variance. Identical observation sequences were presented across subjects. The bottom plots show, for one of the sequences in each task and each observation in the sequence, the estimates of different subjects, their average and the normative estimate. The total deviation (mean squared error) between the subjects’ estimates for a given observation and the normative estimate is equal to the sum of two terms: the bias and the variance, which respectively measure systematic and non-systematic deviations across subjects. We calculated the proportion of bias/variance across sequences. The bar graphs at the top show the total proportion across sequences ± s.e. (obtained by bootstrapping) for the magnitude (A) and probability (B) task.

Using this method, we found that only a small proportion of the deviations were systematic: in total across sequences, the bias/variance ratio was 23.5/76.5% (±1.1% s.e.) in the magnitude learning task and 17.3/82.7% (±0.7% s.e.) in the probability learning task (Fig. 6). This indicates that there are relatively few systematic deviations in average human learning.

We observe that there is variability in subjects’ estimates, and this variability seems to be autocorrelated over time (as seen in the examples in Fig. 6). Such a variability can be produced in a model by introducing noise in the learning process. In other studies, introducing learning noise helped better explain subjects’ choices, and resulted in an adjustment of choice stochasticity (also known as exploration-exploitation tradeoff) to the surprise level (Drugowitsch et al., 2016; Findling et al., 2019). These observations led us to wonder whether such a learning noise could give rise to the adjustments of the learning rate we observed in subjects. We examined this possibility through simulations of learning noise (Fig. S5), with a noise level that could be constant, as in (Drugowitsch et al., 2016), or scaled to the magnitude of the prediction error, as in (Findling et al., 2019). In both cases, these simulations did not reproduce any of the learning rate adjustments observed in subjects (Fig. S5C). Therefore, the results reported in our study regarding the subjects’ ability to dynamically adapt their learning rate are not explained by learning noise.

## Discussion

Many studies in psychology and neuroscience use a fixed learning rate to model the subject’s behavior (with models known by various names such as delta rule, Rescorla-Wagner, temporal difference (TD) learning, Q-learning, SARSA…) (O’Doherty et al., 2003; Rescorla & Wagner, 1972; Rosenblatt, 1961; Schultz et al., 1997; Sutton & Barto, 2018). Our results actually show that in humans, the learning rate varies dynamically from one observation to the next based on what is observed, and is normatively adapted in response to inferred changes in the environment (Fig. 2). We found evidence for these normative dynamic adjustments of the learning rate in two common types of learning, namely magnitude learning and probability learning, despite fundamental differences that exist between these two types of learning, demonstrating the generality of dynamic adaptive learning in humans.

The dynamic adaptation we have shown for probability learning is distinct from an average adaptation to global environmental statistics such as volatility. In the latter, the average learning rate was found to be higher in a block of trials containing many change points (volatile block) than in a block containing no change points (stable block) (Behrens et al., 2007; Browning et al., 2015; Cook et al., 2019). Here, the adjustment of the learning rate we report is not on average across trials but from trial to trial, in response to single change points. This dynamic adaptation implies an adaptation to volatility (because more volatile environments have more change points, inducing more frequent increases in the learning rate and thus a higher average learning rate), but the reverse is not necessarily true. For instance, adaptation to volatility can arise from a switch between strategies that are tailored to different volatility conditions, with each strategy having its own static learning rate; for example, in a choice task, a strategy closer to choice repetition can be used in the stable condition and one closer to win-stay, lose-shift in the volatile condition (Cook et al., 2019).

The generality of our finding is further strengthened by several advantages of our paradigm compared to the numerous probability learning paradigms based on choices used previously, such as probabilistic reversal learning paradigms (Behrens et al., 2007; Browning et al., 2015; Cook et al., 2019; Cools et al., 2002). First, our measure of the subjects’ learning rate does not require any model. It is obtained directly from the subjects’ behavior (equation [1]). It is therefore not sensitive to the researcher’s modeling assumptions, as is the case when the learning rate is estimated by fitting a model parameter across trials. Second, change points in our tasks are not limited to simple reversals and the times at which they occur are completely unpredictable). This eliminates the possibility that subjects may have anticipated the change points (as do some models of reversal tasks (Costa et al., 2015; Wilson et al., 2014)).

Before further discussing the implications of our findings, we would like to highlight the context in which they were obtained. Here, we aimed to study how subjects learn the hidden variable (a magnitude or a probability) of a given generative process, not how they learn the structure of a generative process. Therefore, we provided subjects with detailed instructions so that they fully understood the generative process of the task, and the goal of estimating a time-varying variable. Learning in a context where the structure is unknown, or partially known (to the extent of the instructions), is a different problem and subject of study known as structure learning (Gershman & Niv, 2010). It requires a distinct theoretical treatment and needs its own empirical investigation. The extent to which our findings apply to structure learning remains to be determined.

Given that dynamic adaptive learning has been demonstrated in our study, research in neuroscience should now uncover its neural mechanisms. To study this question, our paradigm could be reused in neuroimaging experiments, and as a starting point, neural measurements could be related to the computational determinants we presented. Catecholaminergic neuromodulators (noradrenaline and dopamine) are good candidates for investigation as several theoretical and empirical works indicate that they may play a role in regulating learning based on surprise and uncertainty (Aston-Jones & Cohen, 2005; Yu & Dayan, 2005). These neuromodulators provide a biological implementation of gating, also known as gain modulation, a computational mechanism that we have previously shown to play a crucial role for artificial neural networks to achieve the very same dynamic adaptations that we have found here in humans (Foucault & Meyniel, 2021).

### Differences between magnitude and probability learning

In light of the differential effects we found in the magnitude and probability learning tasks (Fig. 5), it will be important in future studies of learning to distinguish the contribution of, on the one hand, the effects related to the immediately received observation and the quantities associated with it, such as error magnitude, surprise and change-point probability, and on the other hand, the effects of prior uncertainty arising from past observations. Moreover, it will be important to distinguish the role of these two types of effects in magnitude learning vs. probability learning, as they do not necessarily involve the same mechanisms in these two types of learning. In the present study, we found that there was overall little correlation between the magnitude and probability task across subjects in terms of how much they weighted change-point probability (no correlation, *r*=-0.01) and prior uncertainty (significant but weak correlation, *r*=0.26), compared to e.g. how frequently they updated their reports (*r*=0.63) (see Supplementary Text 4 for more details). This invites us to be particularly careful and to not consider a priori that the results discovered in one type of learning will generalize to all types of learning under uncertainty, because it runs the risk of confusing what are potentially very different processes.

Comparing previous fMRI studies conducted separately on magnitude learning (McGuire et al., 2014) and probability learning (Bounmy et al., 2023; Meyniel & Dehaene, 2017), we noted differences between the two in the neural correlates that were specific to each type of effect. Specifically, when examining the selective effects of the error (quantified by change-point probability or surprise), we noted that frontal regions were implicated mainly in probability learning (right superior frontal gyrus, left and right supplementary and cingulate eye fields). In magnitude learning, they involved primarily medial regions of the visual cortex (calcarine sulci), whereas in probability learning the effects in visual cortex were more lateral (left/right V3/V4). When examining the selective effects of prior uncertainty (quantified by relative uncertainty or confidence), we noted that ventromedial and anterior prefrontal cortex regions were involved primarily in magnitude learning. Interestingly, both types of learning showed neural correlates in the parietal cortex, but in magnitude learning these correlates were specific to prior uncertainty, whereas in probability learning they were common to both prior uncertainty and surprise.

In general, several factors may give rise to differences between the two types of learning. First, there may be distinct neural mechanisms for processing different types of quantities (magnitudes vs. probabilities). Second, the ranges and dynamics of change-point probability and prior uncertainty are different in magnitude and probability learning (Fig. 4), and different neural substrates may be recruited in these different regimes. Third, these two factors may be processed differently in the two types of learning because, in magnitude learning, change-point probability is more diagnostic than prior uncertainty for detecting change points, while in probability learning, prior uncertainty is a more reliable signal for inferring a change point in the recent past (Fig. 4 E and F). In addition, there may be a trade-off in the degree of processing one can allocate to compute and use these two factors that favors the most relevant factor in a given learning context, namely the change-point probability when learning magnitudes and the prior uncertainty when learning probabilities (Fig. 4A, 5 A and B).

### Frequent updating during the learning process

A previous study proposed that the brain does not update its probability estimate at each observation, but only on rare occasions (Gallistel et al., 2014). This conclusion was based on an empirical observation: in Gallistel et al.’s study, subjects overtly updated their reported probability estimate only once every 18.1 observations on average, with only 7% of updates occurring after a single observation. However, a lack of updating in the subject’s report may not necessarily reflect a lack of updating in the subject’s internal learning process. In contrast to Gallistel et al.’s study, our study shows frequent updating during the learning process. Subjects updated their estimate at each observation in the vast majority of cases: 84% of updates occurred after a single observation (vs. 7% in Gallistel et al.), and on average subjects updated their estimate every 1.4 observation (vs. 18.1) (Fig. S3). This makes our data on human probability learning rather novel.

This difference in the frequency of report updates between Gallistel et al.’s study and our study can be explained by task differences: the cost for the subject to overtly update their estimate greatly varies depending on how the task is designed, and when this cost is high, subjects may not manifest their update. In Gallistel et al.’s task, the action required to update the report induces a time cost which, by reproducing the task interface, we estimated to be 4 to 5 s per observation. If subjects had performed this action for all 1,000 observations of a session, it would have taken them more than an hour (estimated 67 to 83 min) per session, instead of the 25.6 min they took on average (Gallistel et al., 2014). In our task, there is no time cost since the observations occur at fixed time intervals. This design choice seems to have made subjects much more willing to express their update. Subjects in our study also received a performance bonus encouraging them to report their updates.

The frequency of subjects’ updating is critical to the results we have shown on the dynamics of the learning rate at the scale of an observation. Note that all our conclusions hold after stringently excluding from analysis all trials in which the report was not updated (Fig. S4).

### Computational models of human learning

Although the human learning algorithm remains to be determined, the results shown here impose strong computational constraints on this algorithm. Models of this algorithm should be able to exhibit the same dynamic adaptations as humans do in response to change points and depending on change-point probability and prior uncertainty (Fig. 2 and 5). They should also be able to reproduce the variability of human learning, and the fact that it is on average close to normative (Fig. 6).

Various models have been proposed in the literature, including the Kalman filter (Kalman, 1960), the Hierarchical Gaussian Filter (HGF) (Mathys et al., 2011), a model estimating the volatility and stochasticity of the environment (assumed to follow a random walk as in the Kalman filter) (Piray & Daw, 2021), the proportional-integral-derivative (PID) controller (Ritz et al., 2018), an adaptive mixture of delta-rules (Wilson et al., 2013), a model with metaplastic synapses guided by a change detection system (Iigaya, 2016), the normative model we used here (which computes the optimal solution via Bayesian inference) (Adams & MacKay, 2007; Behrens et al., 2007; Heilbron & Meyniel, 2019), particle filters (which uses sampling to approximate the optimal solution) (Brown & Steyvers, 2009; Prat-Carrabin et al., 2021), and several architectures of recurrent neural networks (RNNs) (Foucault & Meyniel, 2021; Wang et al., 2018). The Kalman filter is a particular case because it qualifies for level 1 but not level 2 of adaptive learning as we presented in the introduction: its learning rate depends on the global level of stochasticity and volatility in the environment, but does not depend on what is observed (the changes in learning rate over time are simply dictated by the number of observations). The other models qualify for level 2 (they dynamically adapt their learning rate), but it remains to be evaluated to what extent they can, in both types of learning, conform to the specific normative properties demonstrated here in humans.

In a previous article (Foucault & Meyniel, 2021), we have shown that a small recurrent neural network (as small as three recurrent units) could solve the probability learning task quasi-optimally and reproduce all the above-mentioned normative properties (Fig. 2 to 5). This model thus provides a low-cost, neurally feasible solution. In addition, it has the advantage of not incorporating a generative model of the environment a priori: through mere exposure to the environment, the network spontaneously develops its dynamic adaptive learning capabilities (which is sometimes referred to as ‘meta-learning’) (Foucault & Meyniel, 2021; Wang et al., 2018). One mechanistic insight from our study was that small networks demonstrated optimal adaptive learning capabilities only when equipped with a gating mechanism. This is interesting to put in perspective with the other proposed models as they often involve multiplicative interactions (i.e. multiplication between two state variables or a state variable and an input variable) like gating.

Using our behavioral dataset, future studies will be able to evaluate and compare models to probe the algorithms underlying human learning. New data can also be collected easily and in large amounts using the experimental paradigm that we make available to the community (1 report per observation every 1.5 s).

### Relevance for artificial intelligence

Using a dynamic adaptive learning rate is relevant for machine learning and artificial intelligence. The two determinants we identified could guide the search for new efficient, adaptive learning methods. One advantage of change-point probability and prior uncertainty, compared to state of the art methods such as Adam, AdaGrad and RMSprop (Goodfellow et al., 2016), is that they do not require to evaluate the cost function of the task, which in many cases is not accessible at the level of one input sample (in our task for example, the cost function is the error between the subject’s estimate and the true quantity, which is hidden). This is especially relevant for learning from few input samples, as in one-shot or few-shot learning. According to our findings, change-point probability might be particularly relevant for one-shot learning in contexts where individual samples are highly informative, and prior uncertainty for few-shot learning in contexts where individual samples are less informative. This could be linked to current research in machine learning aimed at improving systems’ ability to assess their uncertainty (Gal & Ghahramani, 2016; Hüllermeier & Waegeman, 2021; Kendall & Gal, 2017; Kompa et al., 2021).

## Methods

The code of the experiment, the collected data, and the analysis code are all available on our GitHub repository at https://github.com/cedricfoucault/ada-learn. This repository allows other researchers to reproduce all the results of the article, and enables them to build upon the materials we have created, including tasks, datasets, and analyses.

### Participants

96 human subjects participated in our study (38 female, median age 30 years, interquartile range 25–37 years, from 19 different nationalities). Participants were recruited from Prolific, an online platform for recruiting participants focusing on academic research. The study was approved by the Comité d’Ethique de la Recherche of the Paris Saclay university (#CER-Paris-Saclay-2023-010). Participants were required to use a computer with a touchpad or mouse and a screen large enough to perform the task. They were rewarded £8.40 for their participation (£9/h at an a priori estimated completion time of 56 min, which also turned out to be the median completion time of the participants), plus a performance-based bonus of up to £4.20 (50% of the base pay). Informed consent was collected before the start of the experiment. No participants were excluded from the analyses.

### Ethics Statement

The study has been approved by the Ethics Committee of the Paris-Saclay University (Comité d’Ethique de la Recherche, approval #CER-Paris-Saclay-2023-010). Participants gave their written informed consent prior to participating in the study.

### Experiment steps

The experiment consisted of the following steps: Task 1 instructions, Task 1 sessions, Task 2 instructions, Task 2 sessions. Each task session consisted of a sequence of 75 observations, taking under 2 min to perform (see examples Fig. 1 E and F). In total, the subjects took a median time of 56 min to complete the whole experiment, including instructions (interquartile range 52–65 min). They performed twenty-one task sessions (six of the magnitude task and fifteen of the probability task; this split was determined a priori with an independent pilot study of eight subjects in order to obtain a sufficiently reliable measure of a subject’s learning rate adjustments, with approximately equal relative error in the measurement in both tasks). At the end of each task session, subjects received feedback whose purpose was to maximize subjects’ engagement with the task. This feedback did not seem to induce a training effect, as performance was stable over the course of the task (Fig. S6). The feedback display showed the subject’s score for the session and the corresponding monetary gain (proportional to the score), and also showed the true values of magnitude/probability along with the subject’s estimates. The score was calculated based on the mean absolute error between subjects’ estimates and the true values of magnitude/probability. The function mapping the error to the score was designed to: normalize the score in 0–100%; keep the score of the normative learner at a constant level (close to 100%); prevent too low scores (equal or close to 0%) that would demotivate subjects, using a softplus nonlinearity.

### Task design

The graphic elements of the tasks were designed with several goals in mind: a) to allow subjects to easily monitor the stimuli as they are appearing and adjust their estimate at the same time, b) to strike a good balance between speed and accuracy of adjustments, c) to share as much as possible between the two tasks (see Fig. 1 A and B and links below to run the tasks). The total length of the slider track was 640 CSS pixels, such that the leftmost and rightmost reachable position of the slider was at about 6–7° of visual angle from the center (1 CSS px corresponds to 0.0213° visual angle at the typical viewing distance of the user’s display, https://www.w3.org/TR/css-values-3/#reference-pixel). This allowed the subject to see the slider and the stimuli at the same time. Tick marks were displayed at every 10% length of the slider to make location easier for subjects. These elements were shared between the two tasks. Additionally, there were a few design elements specific to the probability learning task: labels were put below the tick marks showing the percentage corresponding to the estimated probability (in the magnitude task, this is irrelevant as the position itself is the estimate); the portions of the slider between 0–10% and 90–100% were hatched to indicate that the hidden probability never lies within these intervals; a small yellow wheel and blue wheel were shown at the left and right edge of the slider track, respectively, to indicate the direction of estimation in relation to the two colors; we chose the blue and yellow colors to be easily recognizable by humans even with color-blindness.

Within the 1.5 s interval separating the onset of each observation, the stimulus offset occurred at 1.3s, and in the 1.3–1.5 s inter-stimulus interval, the slider thumb was highlighted (lighter stroke color) to indicate to subjects when their estimate was recorded for calculating their performance.

Throughout the session, subjects could at any time move the slider via motion tracking to update their estimate. Motion tracking was implemented by continuously tracking movements of the subject’s pointer (corresponding to finger movements on a touchpad, or to mouse movements). The pointer itself was hidden during the task, to make it clear to the subject that they were controlling the slider.

In the task instructions, for the magnitude learning task, the choice of the cover story (a hidden person throwing snowballs) was inspired by (Prat-Carrabin et al., 2021).

The tasks can be tested directly online without downloading the repository.

● The magnitude learning task can be tested separately at https://run.pavlovia.org/cedricfoucault/ada-learn-task/ada-pos-study.html?skipConsent
● The probability learning task can be tested separately at https://run.pavlovia.org/cedricfoucault/ada-learn-task/ada-prob-study.html?skipConsent
● The experiment combining both tasks can be tested at https://run.pavlovia.org/cedricfoucault/ada-learn-task/ada-pos-prob-study.html (to perform the magnitude task first) or https://run.pavlovia.org/cedricfoucault/ada-learn-task/ada-prob-pos-study.html (to perform the probability task first).

### Observation sequences and sequence-generating processes of the tasks

We generated a set of 100 sequences for the magnitude learning task and 150 sequences for the probability learning task. Each time a subject performed the tasks, they were shown sequences randomly sampled without replacement from these sets.

For the sake of clarity and to be able to compare the two tasks, we use here as the reference unit the horizontal position within the slider interval normalized between 0 and 1 (left and right edge of the slider track, respectively). This unit applies to the estimates (slider positions) and the hidden magnitudes/probabilities these estimates are about, and to the observations (for the probability task, observation ‘yellow’ and ‘blue’ correspond to ‘0’ and ‘1’, respectively, since when the estimate is equal to 1 this corresponds to a 100% probability of the observation being blue). For the magnitude learning task, the observation-generating process was the same as that used by (Nassar et al., 2012) in order to replicate their procedure: the observations were drawn from a Gaussian distribution with a standard deviation of 10/300 (in normalized unit), and whose mean (the hidden magnitude) changed with a probability of zero for the first three observations following a change point and 1/10 for all trials thereafter. For the probability learning task, the observations were drawn from a Bernoulli distribution whose parameter (the hidden probability) changed with a probability of zero for the first six observations following a change point and 1/20 for all trials thereafter. The probability was sampled uniformly between 0.1 and 0.9 initially, and resampled uniformly in the same interval at each change point subject to the constraint that the resulting change in the odds p/(1-p) be no less than fourfold. These parameters were chosen to maintain continuity with previous studies (Bounmy et al., 2023; Heilbron & Meyniel, 2019; Meyniel et al., 2015; Meyniel & Dehaene, 2017). Sampling probabilities between 0.1 and 0.9 avoids producing sequences with excessively long streaks of identical observations where almost no update needs to be made. The minimum distance between change points and the odds-change constraint avoid having change points that are too short-lived or too subtle to be perceived (even the optimal solution shows almost no learning rate adjustments in response to such changes). Other studies on probability learning also used probabilities between 0.1 and 0.9 and constrained change points such that nearly imperceptible change points did not occur (for comparison, the often-used reversal between 20% and 80% produces an odds change of 16) (Behrens et al., 2007; Browning et al., 2015; Cook et al., 2019).

Note that, for the magnitude learning task, we used the same parameter values as previous studies (Nassar et al., 2012; Vaghi et al., 2017). Equalizing the two tasks in terms of the informative value of the observations would not be practically feasible (see also Fig. S1).

### Behavioral analyses

The subject’s estimate *v_t_*, for observation *x_t_*, was taken as the normalized position of the slider at the end of the 1.5s interval just before receiving the next observation (the estimate prior to the first observation, *v_0_*, is 0.5, as the slider is initially positioned in the middle). The value *x_t_* is equal to, for the magnitude task, the (normalized) position of the stimulus, and for the probability task, 1 for a blue draw and 0 for a yellow draw. *v_t-1_* and *v_t_* are the subject’s estimate prior to and after receiving that observation, and *α_t_*is the subject’s learning rate for that observation.

In the results relating to the subjects’ estimate accuracy, the significance was calculated by computing the Pearson correlation at the subject-level between the subject’s and the normative estimates, and then performing at the group-level a two-tailed, one-sample t-test of those correlations against zero.

When analyzing learning rates in the magnitude learning task, similar to (Nassar et al., 2012; Vaghi et al., 2017), we ignored from the analyses outliers in the learning rates (which might occasionally occur when the error is very close to 0), corresponding to learning rates above 1.3 or below -0.6. These values were determined a priori using the distribution of the subjects’ learning rates in an independent pilot study of eight subjects by taking the values above and below which the empirical density was less than 0.05.

### Normative models

Normative models are Bayesian models that compute the posterior distribution *p(h_t_ | x_1:t_)*, which is the probability distribution over the latent quantity that generates the observations (*h_t_*) given the observations received by the subject (*x_1:t_*), with knowledge of the true generative process of observations of the task (Ma et al., 2023).

Given the properties of the generative process in both tasks, this distribution can be computed sequentially using the following update equation (this method is known as Bayesian filtering, for a textbook, see (Särkkä & Svensson, 2023)).

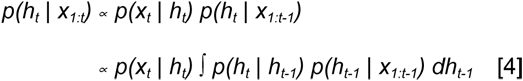

where α denotes equality up to a normalization constant (the left-hand side is obtained from the right-hand side by dividing the right-hand side by its sum over the possible values of h_t+1_, so that the distribution sums to 1). This equation is derived by applying the rules of probability theory and leveraging two conditional independence properties of the generative process: h_t+1_ is conditionally independent of x_1:t_ given h_t_, and x_t+1_ is conditionally independent of x_1:t_ given ht+1.

Having computed the posterior distribution, its mean can be computed to obtain the model’s estimate, *v_t_ = E[h_t_ | x_1:t_]*, and its standard deviation to obtain the uncertainty, which is the prior uncertainty for the next time step (*u_t_*=*SD[h_t-1_ | x_1:t-1_]*, equation [2]). The change-point probability can be computed from the posterior distribution using the following equation, which is derived from Bayes’ rule by leveraging the two conditional independence properties mentioned above:

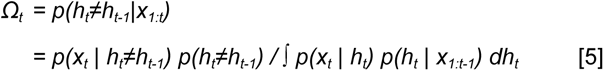

Below, we provide the details of the specific implementation of the model we used for each task.

#### Probability learning task

We reused the implementation of the Bayesian model for the probability learning task of (Meyniel, 2020) whose code is available online at https://github.com/florentmeyniel/TransitionProbModel. We made one modification to match the task asked of the subject in the present study: as the subject was told that the hidden probabilities are drawn in [0.1, 0.9] (with the restriction of the slider to that interval), we also incorporated this knowledge into the model, by setting the corresponding distribution, *p(h_1_)* and *p(h_t+1_ | h_t+1_≠h_t_)*, equal to the uniform distribution on [0.1, 0.9], rather than [0, 1] as in the original version.

#### Magnitude learning task

We implemented the reduced Bayesian model of the magnitude learning task described in (Nassar et al., 2012), which has also been used in other magnitude learning task studies (McGuire et al., 2014; Vaghi et al., 2017). We implemented this reduced model rather than the full model to replicate the procedure of Nassar et al, 2012, whose magnitude learning task serves as a reference for comparison with the probability learning task. This model simplifies computations by only maintaining the first two moments of the posterior distribution, which was reported to have minimal effect on estimates in this task (Nassar et al., 2010, 2012). The model variables are computed sequentially using the following system of equations:

The estimate of the latent mean:

b_t+1_ = b_t_ + η_t_ (x_t_ - b_t_) vt = bt+1

Learning rate:

*η_t_ = τ_t_ + (1-τ_t_) Ω_t_*

The change-point probability:

*Ω_t_ =* 𝒰*(x_t_) H / [*𝒰*(x_t_) H +* 𝒩*(x_t_; b_t_, σ_t_^2^) (1-H)]*

The predictive variance of observations:

*σ_t_ ^2^ = N^2^ + τ_t_ N^2^ / (1 - τ_t_)*

The relative uncertainty about the latent mean with respect to the predictive variance of observations:

*τ_t+1_ = [N^2^ Ω_t_ + (1-Ω_t_) τ_t_ N^2^ + Ω_t_ (1-Ω_t_) (x_t_ τ_t_ + b_t_ (1- τ_t_) - x_t_)^2^] / [N^2^ Ω_t_ + (1-Ω_t_) τ_t_ N^2^ + Ω_t_ (1-Ω_t_) (x_t_ τ_t_ + b_t_ (1- τ_t_) - x_t_)^2^ + N^2^]*

The uncertainty about the latent mean:

*u_t_^2^ = τ_t_ σ_t_^2^*
where *𝒰* is the uniform distribution, *𝒩* is the normal distribution with the mean and variance given after the semicolon, *N* is the standard deviation of the observation generation distribution of the task generative process, and *H=p(h_t_≠h_t-1_)* is the probability of occurrence of a change point of the task generative process.

### Bias/variance decomposition of the mean squared error

Having presented the same sequences multiple times across subjects in our study, we have, for a given sequence *k* out of *K* sequences, *N(k)* subjects who provided estimates (*K*=100 and 150 sequences for the magnitude and probability tasks respectively; median *N(k)* across sequences: 6 and 10 for the magnitude and probability tasks

respectively). For a time step *t* of the sequence, the mean squared error (*mse*) between the normative estimate (*v_t_^n^*) and the subjects’ estimate (*v_t_ ^s(i)^*, where *s(i)* denotes the *i*-th subject among the *N(k)* subjects) is written as:

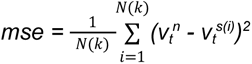

This is equal to the sum of the squared bias error (also called “bias error”) and the variance (also called “variance error”) (Hastie et al., 2009). The squared bias error (*sbe*) is written as:

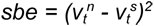

where 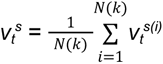 is the mean estimate across subjects.

The variance (*var*) is written as:

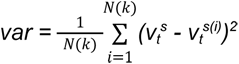

We quantified the precision of our measurement of the proportion of bias/variance (and of the proportion of conservatism bias in Table S3) by the standard error (s.e., which corresponds to the margin of error of the 68% confidence interval of the measurement). To estimate it, we used a bootstrapping procedure. We generated 10,000 new sets of sequences of the same size as the original set by resampling the original set with replacement, and performed the measurement on each of those sets. The estimate of standard error is the standard deviation of the measurement across the sets.

## Data and code availability

All the data and code to reproduce the results of this article are available on our repository at https://github.com/cedricfoucault/ada-learn. This repository also contains the code of the experiments we created, which can be used to collect new data.

## Acknowledgements

We gratefully acknowledge the support received for this work. C.F. was supported by a PhD fellowship from ENS Paris-Saclay (France). F.M. was supported by the European Research Council (ERC grant #947105) and the Inserm. We thank Maëva L’Hôtellier and Alexander Paunov for useful feedback on the project. We also thank them as well as Steven Geysen and Tiffany Bounmy for piloting the experiment.

## Author contributions

CF: Conceptualization, Formal analysis, Investigation, Methodology, Software, Visualization, Writing—original draft preparation, Writing—review & editing; FM: Conceptualization, Funding Acquisition, Methodology, Supervision, Writing—original draft preparation, Writing—review and editing.

## Supplementary information

### Supplementary Text and Tables

#### Supplementary Text 1: Comparison between normative estimates and subjects’ estimates

##### Linear regression

To see how subjects’ estimates compared to normative estimates, we performed a linear regression between the two at the subject level, and then summarized the results at the group level. We collected two measures derived from the regression: the Pearson correlation coefficient and the slope of the linear regression. The results are reported in Tables S1 and S2 below.

The Pearson correlation coefficient results (Table S1) had already been reported in the main text. This coefficient measures the strength of the linear relationship between normative estimates and subjects’ estimates. The fact that the coefficient is significantly greater than 0 shows that subjects’ estimates covary with the optimal estimates, indicating that subjects perform the task adequately.

The slope of the linear regression indicates by how much the subject’s estimate changes on average when the normative estimate changes by one unit. Many studies on human estimates, especially for probability judgments, have observed that the slope of the regression was less than 1 (Costello & Watts, 2014; Erev et al., 1994; Hilbert, 2012; Phillips & Edwards, 1966; Zhu et al., 2020). Consistent with these studies, we also observed in our study that the slope was less than 1, in both tasks (see Table S2 for descriptive and inferential statistics). In the literature, this phenomenon has been referred to as “conservatism bias” (Costello & Watts, 2014; Erev et al., 1994; Hilbert, 2012; Phillips & Edwards, 1966; Zhu et al., 2020), because a regression with a slope less than 1 predicts that, for a given level of normative estimate, the subject’s estimate will be on average less close to the extremes (0 or 1, hence the ‘conservatism’ label), i.e. closer to 0.5, than the normative estimate. Here, we do not attach any particular mechanistic interpretation to the slope and treat it as a descriptive measure. For possible explanations of this phenomenon, see (Costello & Watts, 2014; Erev et al., 1994; Hilbert, 2012; Zhu et al., 2020).

##### Decomposition of the mean squared error

As presented in the main text, we performed a decomposition of the mean squared error between the subjects’ estimates and the normative estimates to quantify the proportion of the error that was attributable to systematic biases in their estimates rather than to their variance (see Results).

We also conducted an additional analysis to investigate the bias: Since we observed a regression slope less than 1 consistent with a “conservatism bias” (Table S2), we investigated the extent to which such a conservatism bias could explain the subjects’ bias. Specifically, we quantified the amount of bias explained by a linear regression model fitted to the subjects, which applies a linear transformation to the normative estimates, and models a conservatism bias when its slope is less than 1. We performed a linear regression between the normative estimates and the subjects’ estimates averaged across the group, took the predictions of this regression as a model of the biased estimates, and then calculated the mean squared error obtained by replacing the normative estimates with the biased estimates. The proportion of the mean squared error that was reduced by using the biased estimates (i.e. the obtained reduction of the error in proportion to the original error) measures the amount of bias explained by the conservatism bias in subjects.

The full results of the decomposition (proportion of bias, variance, and of conservatism bias) are reported in Table S3 below.

#### Supplementary Text 2: Regression on subject’s learning rate in the magnitude learning task performed as in a previous study

For comparison with previous studies on magnitude learning, we additionally performed a regression analysis on the subject’s learning rate as it was done in (McGuire et al., 2014), that is, without z-scoring regressors as we did in the regression reported in the main text, and replacing our prior uncertainty regressor by the RU*(1-CPP) regressor. We obtained regression weights similar to but slightly higher than those reported in (McGuire et al., 2014): The median and interquartile range of the regression weights in this analysis are 0.83 [0.56–0.92] and 0.51 [0.24–0.73] in our data for change-point probability and RU*(1-CPP) respectively (all two-tailed signed-rank p<0.001), vs. 0.53 [0.40–0.76] and 0.32 [0.11–0.44] in (McGuire et al., 2014).

#### Supplementary Text 3: Noisy delta-rule simulations

To examine the possibility that the learning rate adjustments observed in subjects could emerge from learning noise, we conducted simulations of a noisy delta rule model, with noise in the update similar to (Drugowitsch et al., 2016; Findling et al., 2019). The model is described by the following update equation:

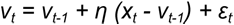

where v_t_ is the model’s estimate following observation x_t_, η is the delta-rule parameter, and ε_t_ is the noise in the update, which is sampled from a zero-mean Gaussian distribution whose standard deviation corresponds to the noise level

We tested two variants for the noise level in the model: one (version a) where, as in (Drugowitsch et al., 2016), it is a constant parameter of the model, *σ_ε_*, and another (version b) where, as in (Findling et al., 2019), it is scaled to the prediction error, with a scaling factor parameter *ζ*. Thus, the noise sampling is *ε_t_ ∼* 𝒩*(0, σ_ε_)* in version (a) and *ε_t_ ∼* 𝒩*(0, ζ |x_t_ - v_t-1_|)* in version (b).

To compute the parameter values, we leveraged the following properties of the model (E denotes the expectation, SD the standard deviation):

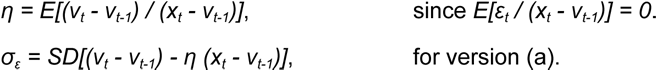

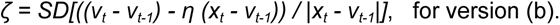

By computing the means and standard deviations described above across time and sequences, we obtained, for each subject and task, the parameter values that best match the subject’s estimates (Fig. S5B).

For the results obtained with this model regarding learning rate adjustments, see Fig. S5C.

#### Supplementary Text 4: Correlations across subjects between the magnitude learning task and the probability learning task

We correlated the regression weights obtained in the magnitude learning task with those obtained in the probability learning task across subjects (the weights were obtained using the same regression as in Fig. 5). The weight of change-point probability was not significantly correlated between the two tasks (*r*=-0.01, p=0.92), and that of prior uncertainty was weakly though significantly correlated (*r*=0.26, p=0.012) (partial Pearson correlation controlling for the individual’s average update frequency, two-tailed p values). This is indeed due to differences between the two tasks: within each task, when performing the same correlation analysis on two halves of the data (even and odd sessions), we obtained strong correlations (these were, in the magnitude and probability task respectively, *r*=0.64 and 0.95 for change-point-probability, *r*=0.38 and 0.74 for prior uncertainty, all p<0.001, two-tailed). For comparison, another behavioral measure, the average update frequency of the subject, was more strongly correlated across subjects between the two tasks: *r*=0.63, p<0.001 (Pearson correlation, two-tailed).

---

**Fig. S1.**
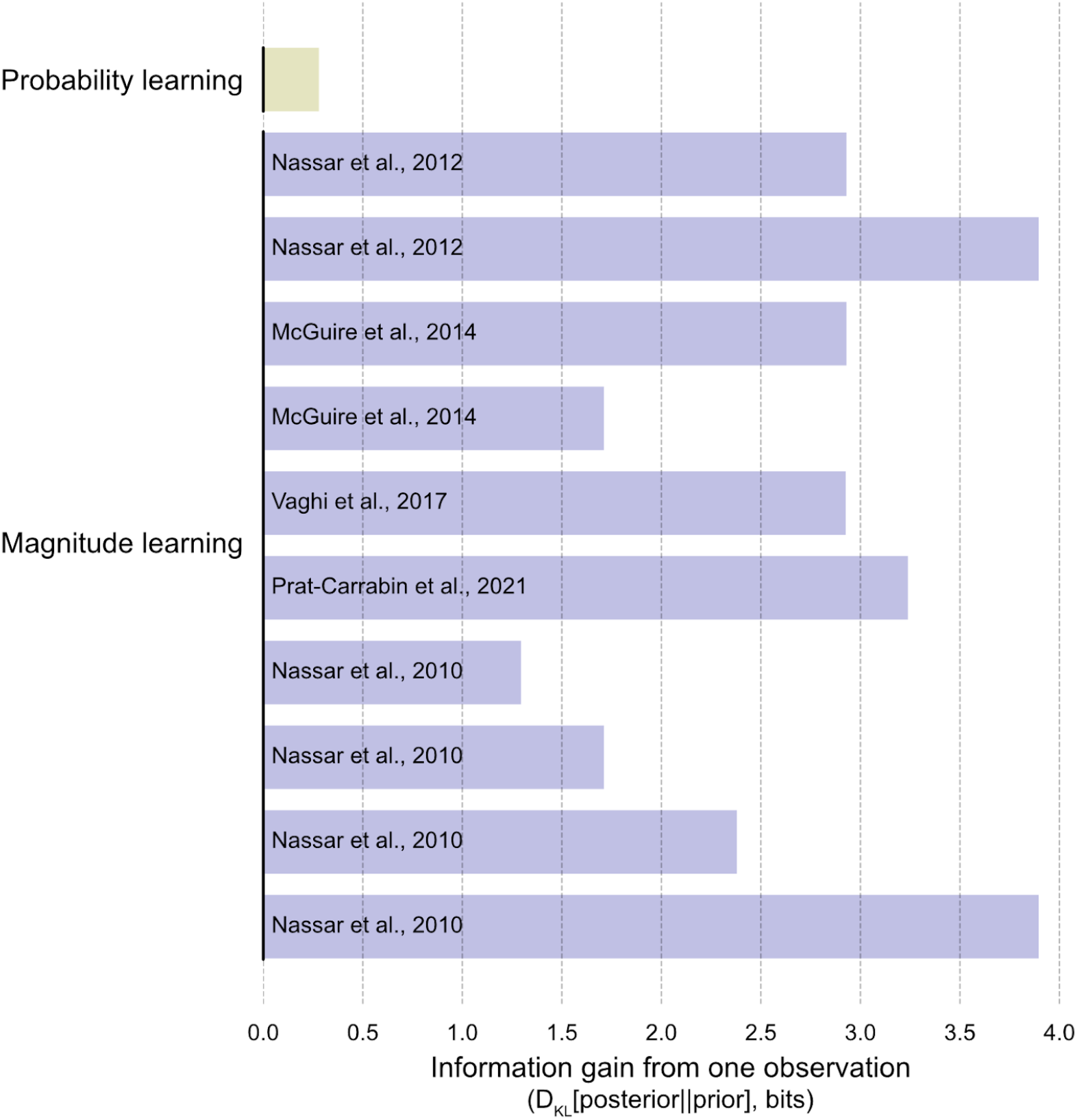
Information gain provided by a single observation about the quantity to be learned in magnitude learning and probability learning. We computed the information gain as the KL divergence between the prior (uniform) distribution before having received any observation, and the posterior distribution about the underlying quantity after having received the observation, using the prior as reference distribution (i.e. D_KL_[posterior||prior]), on average over the possible observations. The posterior is obtained from the prior and the likelihood function relating the observation to the underlying quantity using Bayes rule. In probability learning, the information gain is minimal. This is due to the binary nature of the observation. In magnitude learning, the information gain is larger because the observation is quantitative and typically fairly representative of the underlying magnitude. Although the latter depends on the experimenter’s choice of standard deviation with which observations are generated, we computed the information gain for numerous experiments previously conducted and each condition of these experiments, and as shown above, in all cases it was substantially higher than that obtained in probability learning (McGuire et al., 2014; Nassar et al., 2010, 2012; Prat-Carrabin et al., 2021; Vaghi et al., 2017).

**Fig. S2.**
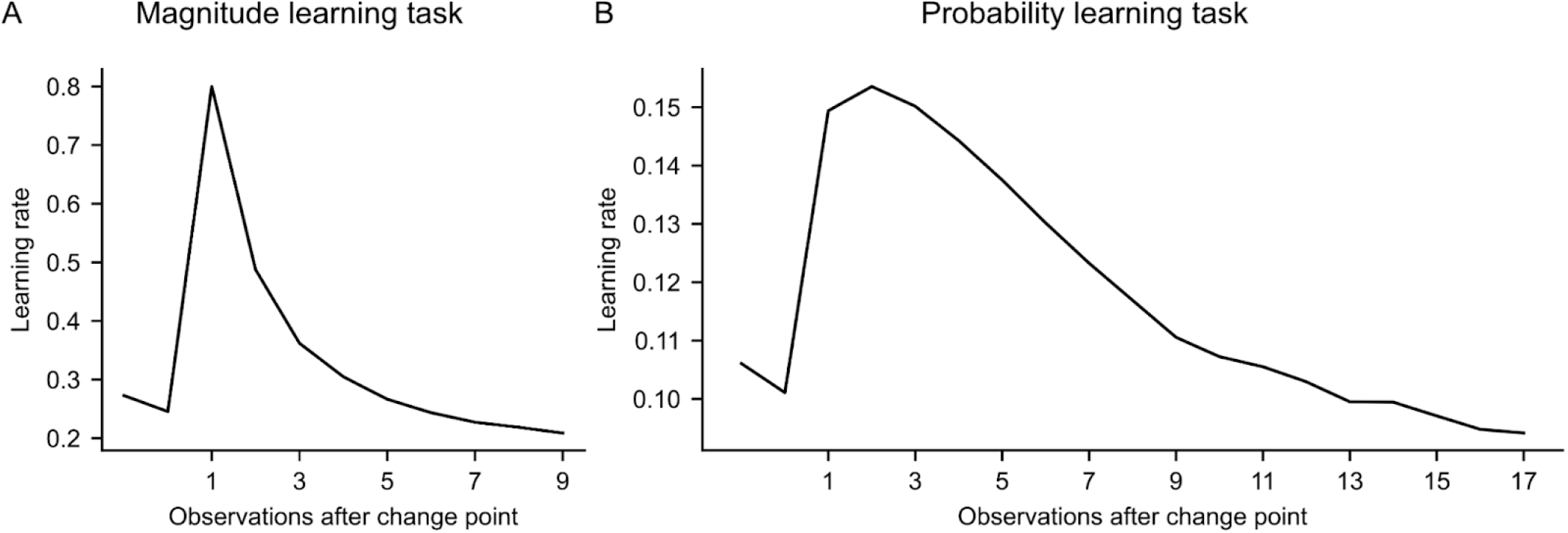
Dynamics of the normative model’s learning rate after a change point, in the magnitude (A) and probability (B) learning tasks. The plots were obtained as in Fig. 2, but rather than using the subjects’ learning rate, we used the normative model’s learning rate instead, which we obtained by running the normative model on the same sequences as the subject.

**Fig. S3.**
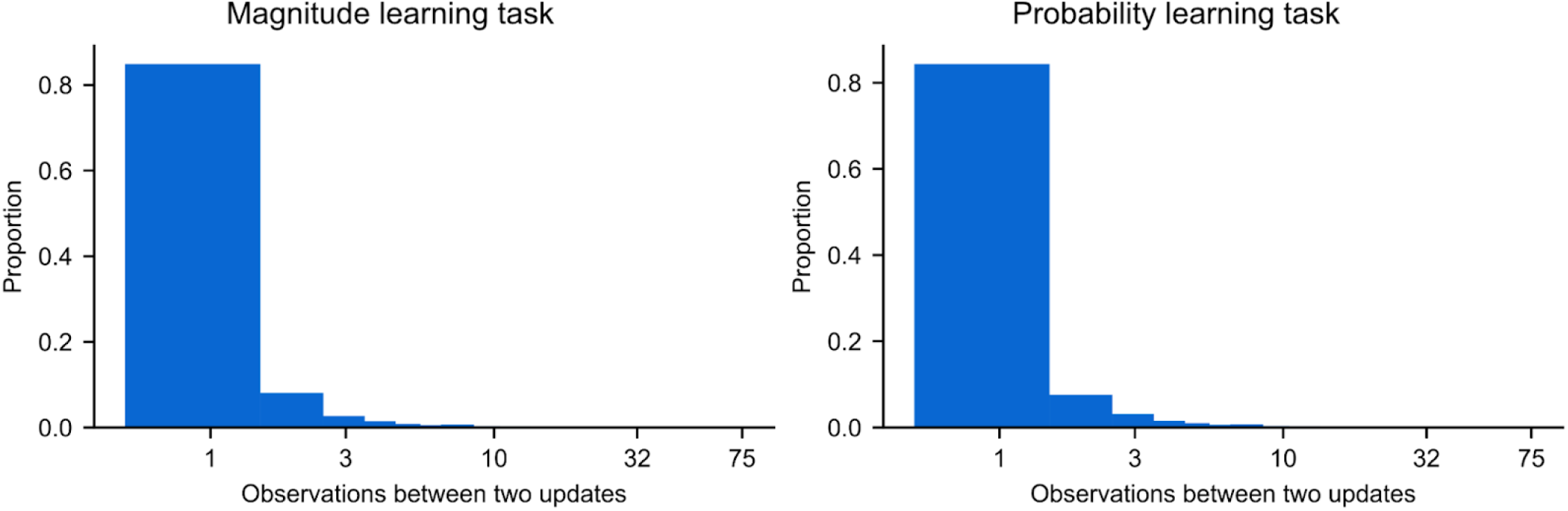
Distribution of the number of observations elapsed between two report updates made by subjects. A log scale was used for the number of observations as in (Gallistel et al., 2014) for comparison (the equivalent distribution in Gallistel et al. is shown in their Fig. 11). In contrast to (Gallistel et al., 2014), in our study, updates were made on each observation most of the time (84% in the above distribution; mean of the distribution: 1.4 observation).

**Fig. S4.**
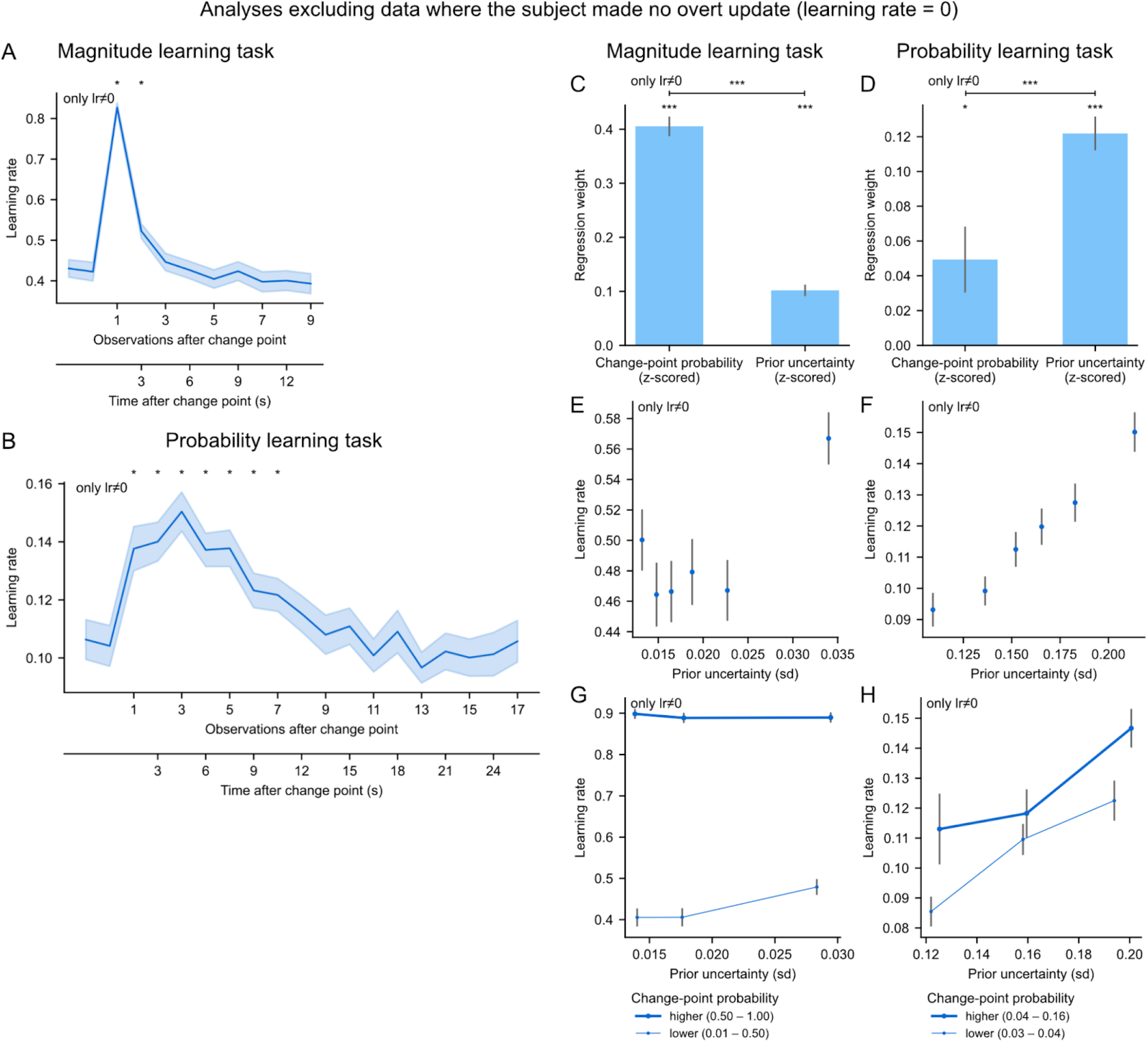
The main results are similar and remain significant when excluding all data where the subject did not make an overt update. After excluding all data points where this was the case (i.e. learning rate = 0), we performed the same analyses as in previous figures and obtained the above plots: (A and B) Equivalent to Fig. 2 A and B; (C–H) Equivalent to Fig. 5 (A–F). Stars denote statistical significance as in the main figures (see legends of those figures for further details).

**Fig. S5.**
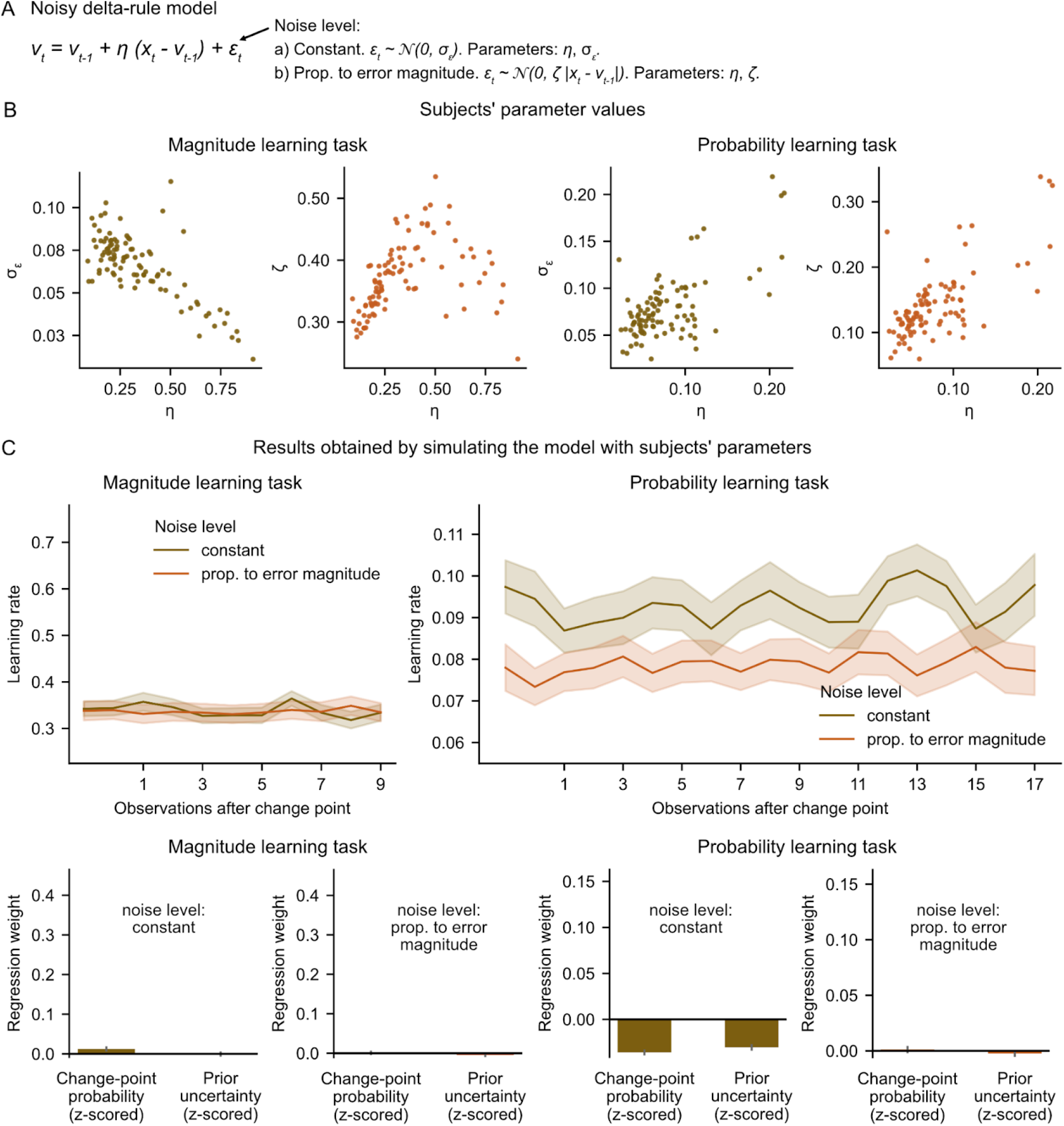
The subjects’ dynamic adjustments of the learning rate are not explained by learning noise. (A) Model of a delta-rule with learning noise. A noise sample is injected at each update of the model, otherwise governed by a delta-rule with parameter η. Two versions were tested for the noise level: (a) constant (parameter σ_ε_), (b) scaled to the magnitude of the prediction error (scaling factor parameter ζ). (B) Values of the model parameters for each subject, for each version of the model and each task. Each dot represents one subject. (C) Results obtained by simulating the model with the subject’s parameters on the subject’s sequences and performing the same learning rate analyses as those reported in the main results for subjects. Top plots are the results for the analysis corresponding to Fig. 2, bottom plots to Fig. 5.

**Fig. S6.**
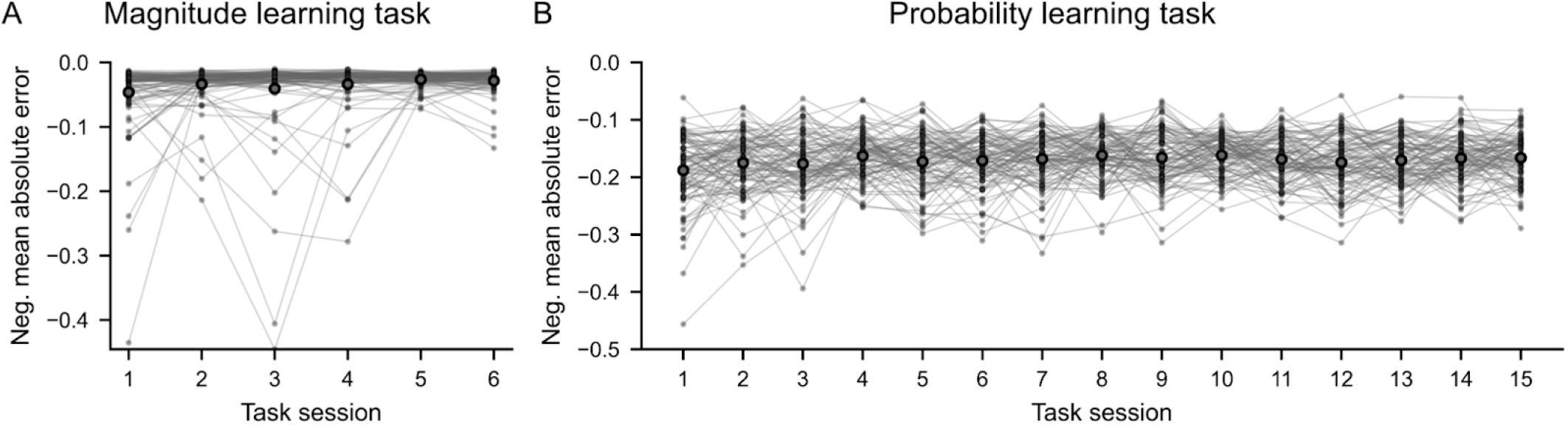
Subjects’ performance was stable over the course of the task. Performance is measured by the accuracy of the estimates, quantified by the mean absolute error between the subject’s estimate and the true value of the hidden quantity (the negative of the error was used so that higher values correspond to higher performance). Thin dots and lines connecting them each denote one subject; large circles denote the mean across subjects.

**Table S1.** Pearson correlation coefficient between normative estimates and subjects’ estimates.

| Task | Mean | S.e.m. | Standard deviation | T-test against 0 |  |
| --- | --- | --- | --- | --- | --- |
|  |  |  |  | t statistic | p value |
| Magnitude learning | 0.96 | 0.01 | 0.09 | 106.20 | 2E-100 |
| Probability learning | 0.80 | 0.01 | 0.14 | 55.79 | 2E-74 |

**Table S2.** Slope of the linear regression between normative estimates and subjects’ estimates.

| Task | Mean | S.e.m. | Standard deviation | T-test against 0 |  | T-test against 1 |  |
| --- | --- | --- | --- | --- | --- | --- | --- |
|  |  |  |  | t statistic | p value | t statistic | p value |
| Magnitude learning | 0.95 | 0.01 | 0.10 | 88.63 | 4E-93 | 4.79 | 6E-06 |
| Probability learning | 0.88 | 0.02 | 0.23 | 37.03 | 3E-58 | 4.96 | 3E-06 |

**Table S3.**
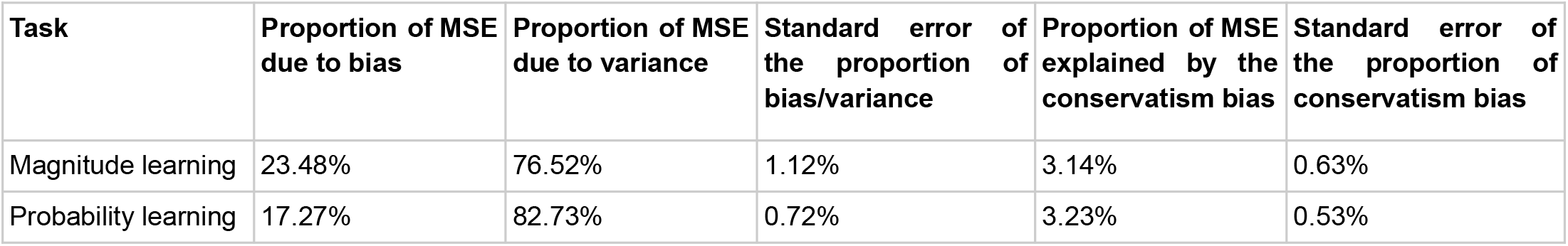
Decomposition of mean squared error between subjects’ estimates and the normative estimates. MSE: mean squared error.

| Task | Proportion of MSE due to bias | Proportion of MSE due to variance | Standard error of the proportion of bias/variance | Proportion of MSE explained by the conservatism bias | Standard error of the proportion of conservatism bias |
| --- | --- | --- | --- | --- | --- |
| Magnitude learning | 23.48% | 76.52% | 1.12% | 3.14% | 0.63% |
| Probability learning | 17.27% | 82.73% | 0.72% | 3.23% | 0.53% |

